# Quadrupling the protein family space with global metagenomics

**DOI:** 10.64898/2025.12.21.695754

**Authors:** Eleni Aplakidou, Fotis A. Baltoumas, Maria N. Chasapi, Efthimia Lamari, IIias Georgakopoulos-Soares, Evangelos Karatzas, Ioannis Iliopoulos, Aydin Buluç, Robert D. Finn, Antonio Pedro Camargo, Nikos C. Kyrpides, Georgios A. Pavlopoulos

## Abstract

The known universe of protein families represents only a small fraction of nature’s molecular diversity. From 40.3 billion sequences across 40,446 metagenomes, 9,540 metatranscriptomes, and 539 million proteins from 167,415 reference genomes, we identified 608,258 previously uncharacterized protein families with ≥100 members and 6.5 million families with ≥25 members, none matching known Pfam domains or reference proteins. This effort doubles the known repertoire of large families and quadruples that of smaller families. Integration of AlphaFold2-based structural predictions with gene-neighborhood and taxonomic analyses enables the characterization of previously unannotated proteins, revealing candidates for both novel and known biological functions in understudied microbial lineages and biomes. This expanded repertoire provides insights into microbial adaptation and broadens the molecular toolkit available for biotechnology, highlighting the power of global metagenomics to uncover hidden protein diversity.

## Main

Proteins serve as the fundamental molecular machines of life, playing essential roles in metabolism, gene expression, cellular defense, and structural integrity. Gaining a comprehensive understanding of proteins is crucial for advancing knowledge in biology, evolution, and disease mechanisms. Although their central role in life, the immense diversity of microbial proteins remains largely uncharacterized, representing a vast reservoir of functional and ecological information yet to be explored. With the explosion of sequencing technologies, we now have access to vast amounts of metagenomic protein sequence data, housed in databases that are growing at an unprecedented rate^1–3^. Existing databases containing protein information, such as sequences, structures, and functional annotations, are confined to well-characterized protein families; others serve mainly as collections of unstructured datasets, and many focus on specific environments, resulting in large portions of protein diversity being overlooked (Supplementary Table 1). Consequently, despite this rapid expansion of data, a vast and largely uncharted metagenomic “protein dark matter” remains underexplored and underrepresented in current repositories. These novel proteins, often hidden within metagenomic datasets, are not merely rare variants; recent studies have shown that they include abundant families that are globally distributed and ecologically significant.

By clustering billions of metagenomic sequences and integrating structural prediction, gene-neighborhood context, and ecological metadata, it becomes possible to define both abundant and rare protein families, identify novel folds, and assign putative functions to previously uncharacterized sequences. This approach not only increases the coverage of microbial protein space but also bridges the gap between sequence discovery and functional insight, enabling targeted exploration of enzymes, metabolic pathways, and molecular scaffolds for biotechnology. In this study, we present a large-scale, rigorously curated expansion of novel microbial protein families. Using a scalable, efficient, biologically validated pipeline on one of the largest microbial datasets assembled to date (Figure 1), we produce a high-quality catalog of novel protein families and folds with potential functions. This framework helps probe the functional dark matter of metagenomes, reveal new structural and biochemical diversity, and enable applications from enzyme discovery to synthetic biology, turning metagenomic sequence space into actionable biological insight.

**Figure 1.**
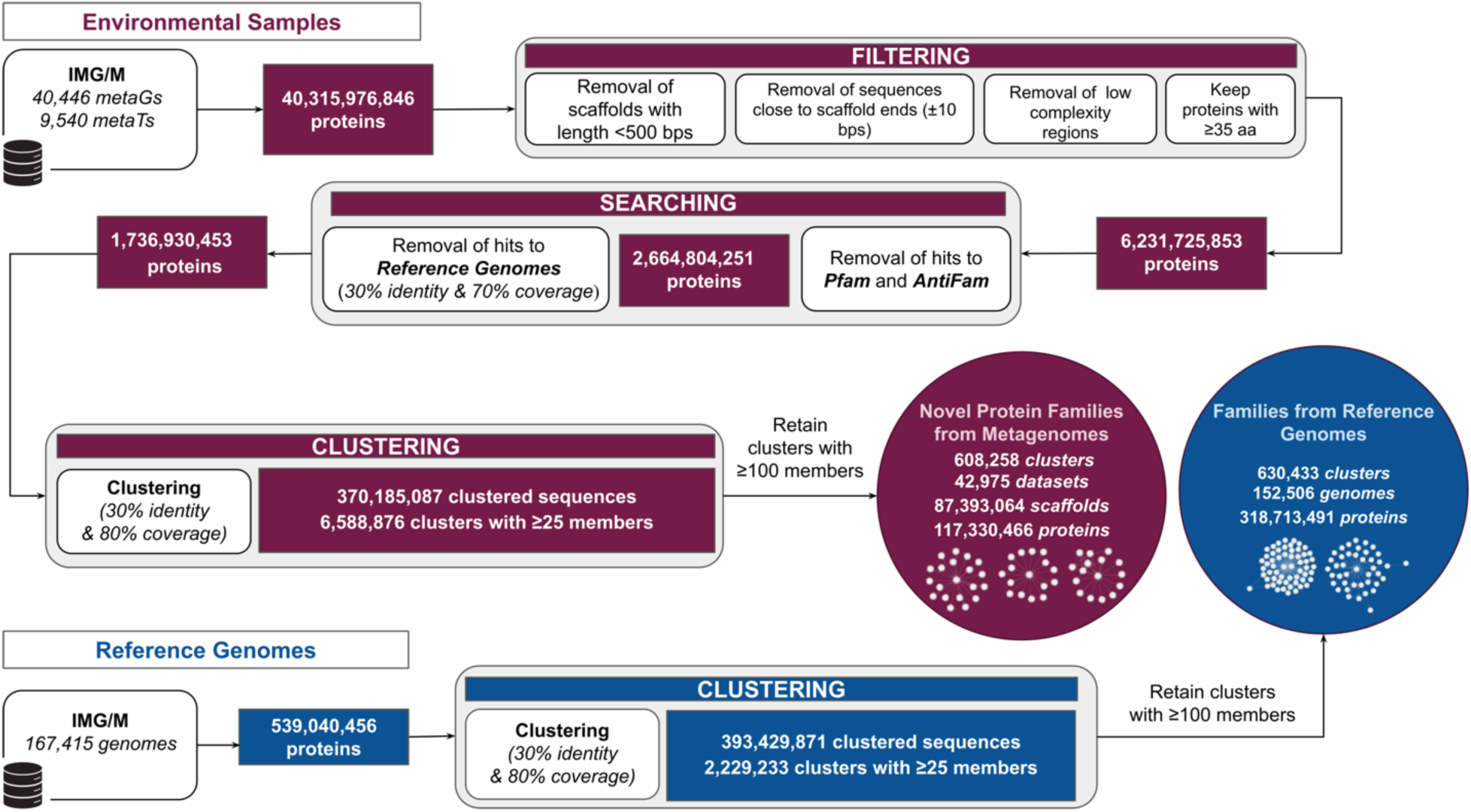
Pipeline illustrating the sequence filtering and clustering methodology for generating protein families from both reference genomes and metagenomes/metatranscriptomes.

## Results

### Novel families from metagenomes vs families from reference genomes

We identified novel protein sequences, the ones with no hits to Pfam^4^, Antifam^5^, or reference genomes. These sequences were subsequently clustered into families using MMseqs2^6^. For direct comparison with known protein families, the same methodology was applied to sequences derived from reference genomes, analyzing them across various cluster sizes (≥25, ≥50, ≥75, and ≥100 members) (Table 1).

**Table 1.** Direct comparison of protein clusters produced from reference genomes versus novel protein clusters generated from metagenomic and metatranscriptomic data at varying cluster sizes (≥25, ≥50, ≥75, and ≥100 members).

| Reference Genomes |  |  |  |  |
| --- | --- | --- | --- | --- |
| Proteins for clustering 539,040,456 |  |  |  |  |
| Family size | $\geq 25$ members | $\geq 50$ members | $\geq 75$ members | $\geq 100$ members |
| Families | 2,229,233 | 1,200,486 | 824,621 | 630,433 |
| Number of Genomes | 155,857 | 154,072 | 153,253 | 152,506 |
| Proteins | 393,429,871 | 358,091,618 | 335,371,741 | 318,713,491 |
| % of proteins | 72.99% | 66.43% | 62.22% | 59.13% |
| Environmental Samples |  |  |  |  |
| Proteins for clustering 1,736,930,453 |  |  |  |  |
| Family size | $\geq 25$ members | $\geq 50$ members | $\geq 75$ members | $\geq 100$ members |
| Families | 6,588,876 | 2,160,632 | 1,052,590 | 608,258 |
| Number of Metagenomes | 44,198 | 43,701 | 43,315 | 42,975 |
| Scaffolds | 253,359,325 | 157,845,393 | 113,497,493 | 87,393,064 |
| Proteins | 370,185,087 | 221,646,880 | 155,259,616 | 117,330,466 |
| % of proteins | 21.31% | 12.76% | 8.94% | 6.76% |
| **Comparison with reference genomes** |  |  |  |  |
| **% of family increase** | **296%** | **180%** | **128%** | **96%** |
| % of protein increase | 94% | 62% | 46% | 37% |

Clustering analysis of reference genomes grouped 393,429,871 of 539,040,456 proteins (72%) into 2,229,233 clusters of ≥25 members across 155,857 genomes. In contrast, metagenomic and metatranscriptomic data clustered 370,185,087 of 1,736,930,453 proteins (21.3%) into 6,588,876 clusters from 44,198 datasets and 253,359,325 scaffolds, representing a 296% increase in protein families and quadrupling the total for this size threshold. At ≥50 members, 358,091,618 proteins (66.4%) from reference genomes formed 1,200,486 clusters, while 221,646,880 proteins (12.8%) from metagenomes grouped into 2,160,632 clusters, reflecting a 62% increase in clustered proteins and a 180% rise in protein families. For ≥75 members, 335,371,741 reference proteins (62.2%) formed 824,621 clusters, versus 155,259,616 metagenomic proteins (8.9%) in 1,052,590 clusters, showing a 46% increase in clustered proteins and 128% more families. At ≥100 members, reference genomes contained 318,713,491 proteins (59.1%) in 630,433 clusters, while metagenomes grouped 117,330,466 proteins (6.8%) into 608,258 clusters, doubling the protein family space.

As clustering thresholds increased, reference genomes showed a minimal reduction in the number of clustered proteins but a notable decrease in the number of clusters, indicating more consolidated protein families. By contrast, metagenomic data exhibited a 30% reduction in proteins within larger clusters and a 50% drop in cluster count, suggesting that metagenomic data tend to form smaller, more fragmented clusters due to greater sequencing fragmentation. Further family properties of protein clusters from reference genomes and metagenomes are shown in Supplementary File 1 and Supplementary Figure 1.

### Biome distribution

Metagenomic and metatranscriptomic datasets from IMG/M^7^ included 49,986 samples with geographic coordinates (latitude and longitude) and GOLD^8^ ecosystem classifications (Environmental, Engineered, Host-Associated). Most samples were Environmental (67.7%), followed by Host-Associated (23%) and Engineered (9.2%). Focusing on microbial families with at least 100 members and at least 5% of samples originating from a single ecosystem type, we identified 608,258 families across 42,975 datasets: 67.5% Environmental, 22.6% Host-Associated, and 9.7% Engineered. Geographic coordinates for Environmental samples were used to plot their origin locations on a world map (Figure 2A), underscoring the global reach and diversity of sampled environments.

**Figure 2.**
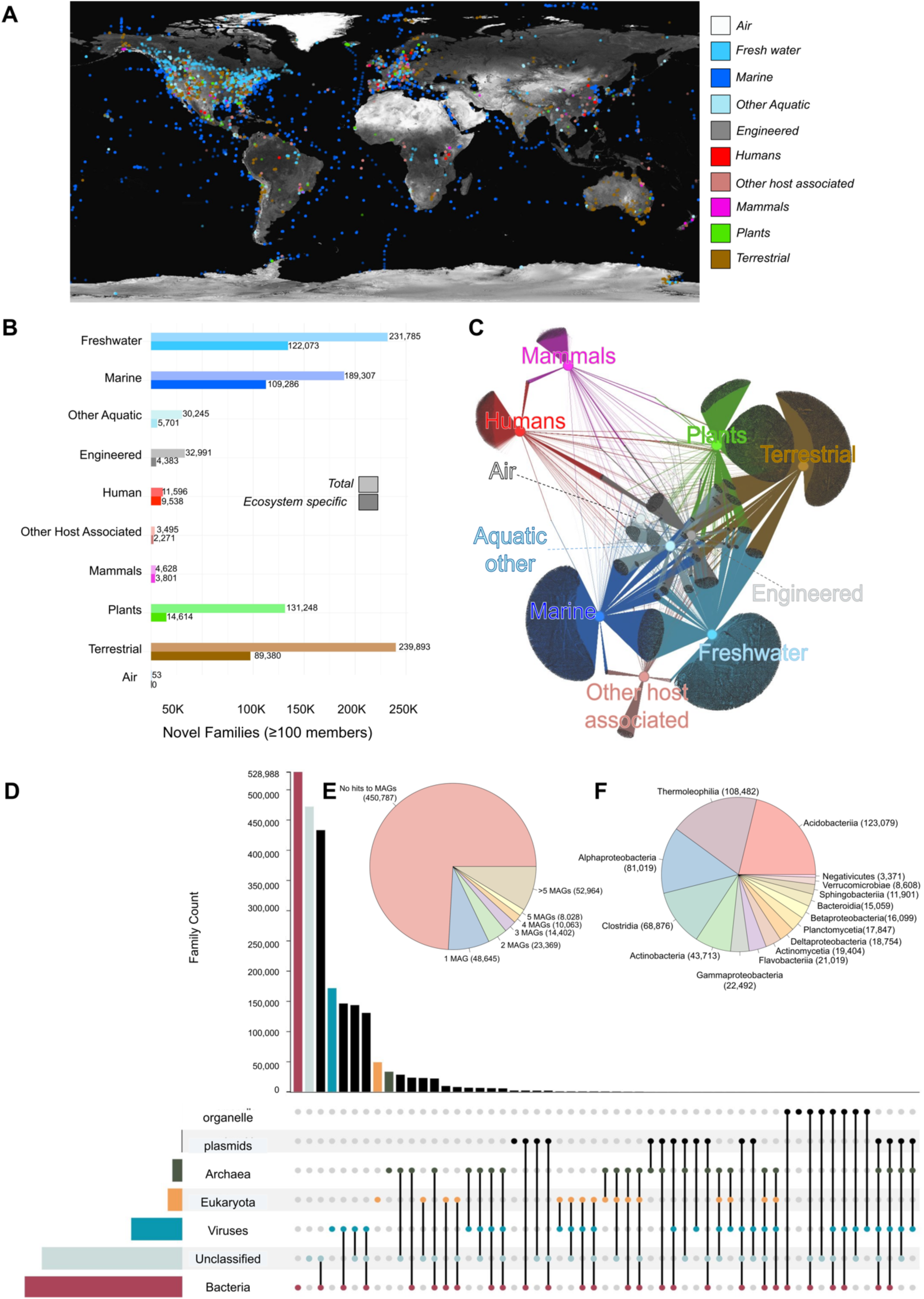
Ecosystem Analysis of Novel Protein Families with ≥100 members (A) Visualization of sample geographical distribution. (B) Network representation of the protein clusters and their ecosystems. Ten ecosystem types were applied according to the GOLD ecosystem classification, represented by central, colored nodes (hubs), whereas the grey peripheral nodes represent the protein clusters. The edges represent the protein cluster–ecosystem associations. (C) The distribution of total versus ecosystem-type-specific NMPFs across the eight different ecosystem types. (D) Family taxonomy distribution across the three domains of life, viruses, plasmids, and organelles. (E) Distribution of protein hits against High-Quality Metagenome Assembled Genomes (HQ MAGs) within families. (F) The top 20 MAGs phyla that are represented across the families.

Of these families, 385,791 (63.4%) are restricted to a single ecosystem: primarily Environmental (57.5%), with smaller contributions from Host-Associated (4.9%) and Engineered (0.7%). Among these ecosystems, families exclusive to a single ecosystem include 89,380 (14.6%) in Terrestrial, 122,073 (20%) in Freshwater, 109,286 (17.9%) in Marine, 5,701 (0.93%) in Other Aquatic Environments, 4,383 (0.7%) in Engineered, 14,614 (2.4%) in Plants, 3,801 (0.62%) in Mammals, 9,538 (1.56%) in Humans, and 2,271 (0.37%) in other host subtypes (Figure 2B). Biome-specific novel protein families highlight previously unrecognized functions and sequence traits that may be adapted to their particular ecological niches (see Gene Neighborhood).

Among families spanning more than one ecosystem, 185,994 (30.5%) exist in two ecosystems. Notable overlaps include 96,049 families (16%) shared between Terrestrial and Plants, and 50,232 families (8.2%) shared between Freshwater and Marine environments. Additionally, 9,539 families (1,5%) are found in Terrestrial and Freshwater ecosystems, and 9,200 families (1.5%) span Engineered and Freshwater environments. Functional overlap between environments can uncover common biological roles among diverse microbial communities. An UpSet plot visualizing all possible overlaps in detail is shown in Supplementary Figure 2.

Figure 2C presents a network representation illustrating habitat-specific families and families shared across multiple environments. In this network, gray nodes represent novel protein families with 100 or more members, while colored nodes denote the different habitats.

### Gene Neighborhood

The analysis of gene neighborhoods within novel protein families, particularly those specific to different environments, provides valuable functional insights into microbial activities across various ecosystems. To gain a more accurate understanding of functions, this study focuses on biome-specific families, aiming to identify the most frequent functions and those that are relatively unique compared to the 50 most common functions in each environment. (See Supplementary Figure 3-5 and Supplementary Table 2-4).

Focusing on human-associated specific protein families, the majority of the analyzed samples originate from the digestive system. Gene neighborhood analysis reveals that the MatE^9^ (Multi Antimicrobial Extrusion) family, a member of the multidrug/oligosaccharidyl-lipid/polysaccharide (MOP) flippase superfamily, is significantly more prevalent in this environment. Metagenomic studies indicate that genes related to MatE contribute to antimicrobial resistance in gut microbiota. Additionally, MFS/sugar transport proteins play a crucial role in nutrient acquisition, enabling the efficient transport of diverse sugars and metabolites across microbial membranes, an essential function in the gut’s nutrient-rich but competitive landscape^10^. In plant-specific environments, microbes must adapt to unique ecological and physiological conditions, including interactions with plant hosts. The Plant mobile domain and PPR repeat are particularly relevant, as they play essential roles in RNA processing and gene regulation, often found in plant-associated microbes and organelles^11^. Additionally, the NB-ARC domain^12,13^, commonly linked to resistance genes, contributes to microbial adaptation and defense mechanisms within plant environments, highlighting the intricate microbial-plant interactions.

In freshwater ecosystems, hotspots for viral and phage activity, gene neighborhood analysis highlights the exclusive presence of viral-related proteins such as Caudovirus prohead serine proteases^14^, phage portal protein^15^, and YqaJ-like viral recombinases^16^. These proteins are crucial for viral replication, assembly, and horizontal gene transfer, emphasizing the central role of viruses in shaping microbial communities. Additionally, the microbial reliance on carbohydrate metabolism in freshwater environments is evident in the exclusive presence of proteins such as chitinases, glycosyltransferases, and transglycosylases^17^. These enzymes facilitate the degradation of plant matter, detritus, and chitinous material, supporting nutrient cycling and microbial adaptation in these nutrient-rich but variable habitats.

In terrestrial environments, gene neighborhood analysis identifies proteins with transaminase activity and carboxypeptidase regulatory-like domains as highly prevalent in soil ecosystems. These proteins are essential for amino acid metabolism, contributing to nitrogen cycling^18^ by facilitating the synthesis, degradation, and transformation of amino acids and organic nitrogen compounds. Additionally, the significant presence of the tripartite tricarboxylate transporter (TTT) family receptor in these environments highlights its role in the uptake of organic acids from plant root exudates, decaying organic matter, and microbial metabolites. These organic acids serve as crucial energy sources and biosynthetic precursors, supporting microbial survival and activity in soil ecosystems.

### Taxonomy

Of the 87,393,064 scaffolds analyzed, 65,471,449 (74.92%) were successfully classified taxonomically, while 21,921,615 (25.08%) remained unclassified. Integrating multiple taxonomic algorithms in the analysis pipeline improved classification rates, reducing the proportion of unclassified scaffolds compared to the previous study. Among the classified scaffolds, the majority were categorized as Bacteria (76.01%), followed by Viruses (14%), Eukaryota (7%), and Archaea (3%), with a small fraction classified as organelles and plasmids. The high classification rate achieved through deep classification enables exploration of taxonomic diversity and the association of specific taxa with particular functions.

Using a threshold of at least 95% of samples derived from a single taxonomic domain, 115,377 families (18.9%) were taxonomically assigned to unique domains, while the majority encompassed scaffolds from multiple taxonomic groups. Specifically, 72,590 families (12%) were exclusively bacterial, 23,782 (4%) were solely eukaryotic, 10,547 (2%) were viral, and 3,124 (1%) were archaeal. The majority of families displayed mixed taxonomic compositions, including combinations such as Bacteria and Unclassified (360,610 families, 59.2%), Bacteria and Viruses (105,338 families, 17.3%), and Viruses and Unclassified (30,355 families, 4.99%) (Figure 2D).

Proteins from families that matched the 21,817 high-quality MAGs were used for taxonomic analysis. Since most MAGs are taxonomically classified, their taxonomy provides valuable insights into family taxonomy. Most families (74%) show no hits against MAGs, while the remaining families exhibit hits in descending order relative to the number of MAGs, indicating that more families have hits in fewer MAGs, while the number of families decreases as the number of MAGs with hits increases (Figure 2E). Of the MAGs, 90% are bacterial, 6% remain unclassified, and 3% are archaeal. A taxonomic analysis of families based on MAGs identifies the top 20 bacterial species most commonly represented across these families (Figure 2F).

### Structural distribution

Focusing on protein families with more than 100 members, 3D structures were predicted using AlphaFold2/ColabFold^19,20^, following the workflow presented in Figure 3A. Of 608,258 families, structural models were successfully generated for 156,609 families. For each family, the model with the highest average pLDDT score was retained. Among these, 26,767 (17.09%) models had a pTM score > 0.7, indicating high-quality models, while 47,823 (30.54%) had a pTM score between 0.5 and 0.7, indicating medium-quality models. This part of the analysis focuses on the 74,590 family models that fall into these two quality categories.

**Figure 3.**
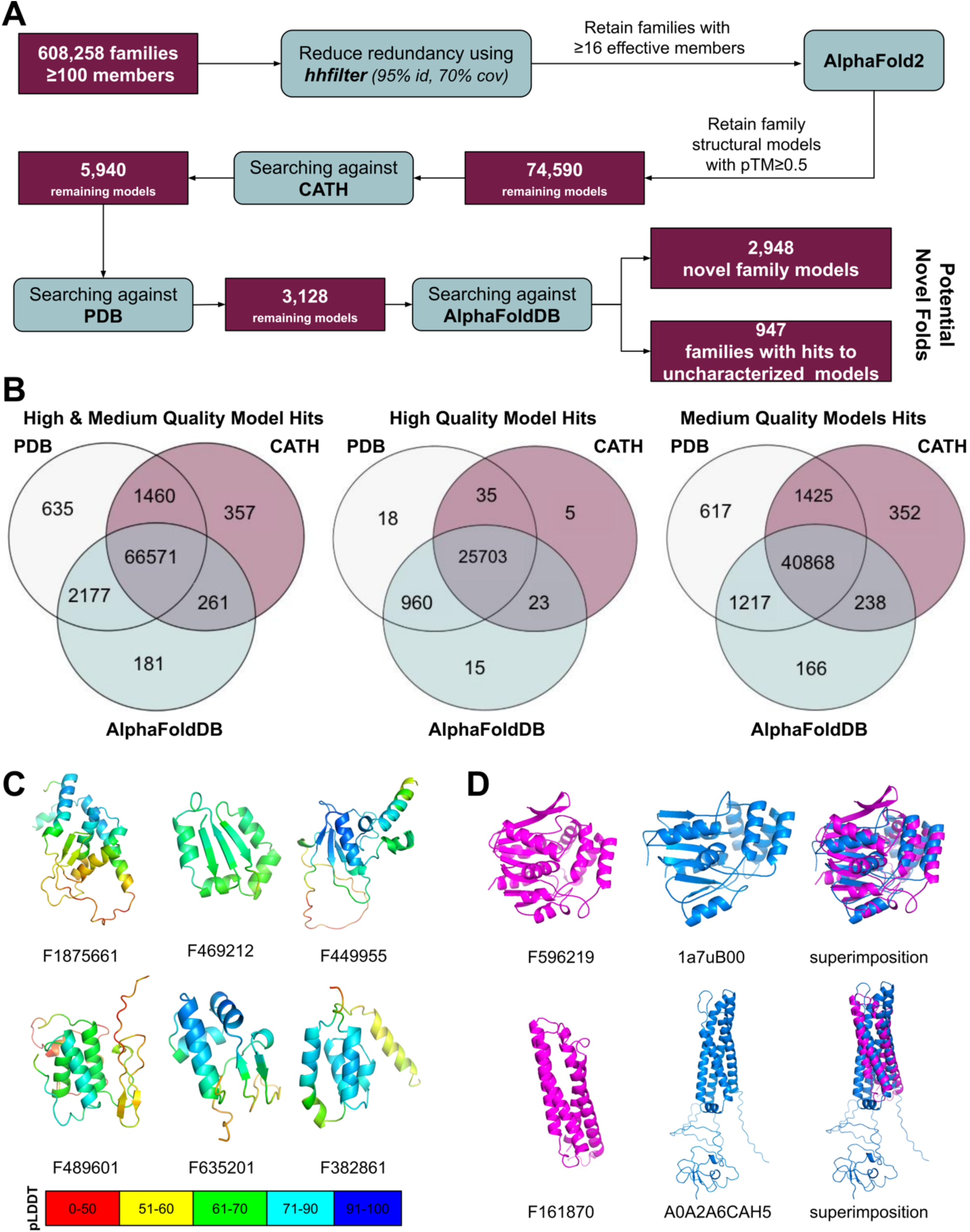
(A) Pipeline illustrating the preparation, structural prediction, and structural search using Foldseek against CATH, PDB, and AlphaFoldDB, highlighting the number of potential novel folds and unannotated structural hits. (B) Venn diagrams for high (pTM≥0.7), medium (0.5≤pTM>0.7), and combined quality model structural hits against CATH, PDB, and AlphaFoldDB, highlighting the overlapping and unique hits for each database. (C) Example models with no significant hits to CATH, PDB, or AlphaFoldDB, considered as potential novel folds. pLDDT, per-residue confidence score. (D) Example models with hits to PDB and AlphaFoldDB domains.

To provide insights into functional annotation and to explore the novel structural space, these models were searched against well-known structural databases that contain both experimentally validated and computationally predicted 3D models. Searches were conducted against CATH^21^, PDB^22^, and AlphaFoldDB^23^ (Figure 3B). The total number of hits for the 74,590 family models included 68,649 hits from CATH, 70,843 from PDB, and 69,190 from AlphaFoldDB. Specifically, searching against CATH yielded 68,649 family model hits, with 25,766 originating from high-quality models and 42,883 from medium-quality models. The remaining unmatched structures were then searched against the PDB, yielding 2,812 additional hits (978 high-quality models, 1,834 medium-quality models). Finally, a search against AlphaFoldDB identified 180 more hits (15 high-quality, 165 medium-quality). An example of structural annotated folds is presented in Figure 3D. Clustering with Foldseek produced 2,781 singletons and 69 multi-member clusters, indicating most folds are unique. Further structural clustering of the whole dataset suggests these models represent genuinely novel partial or complete folds, rather than variations of known architectures.

These findings enhance functional annotation by leveraging structurally characterized proteins and uncover 2,948 potential novel folds (Figure 3C). Notably, 947 family models exhibit structural similarities to entries in AlphaFoldDB, yet their corresponding structures remain uncharacterized. This not only expands the pool of potential novel folds, as many remain unidentified, but also presents an opportunity to infer functional annotations for these previously uncharacterized structures.

### Functional annotation through structural modeling

Structural annotations were performed based on structural homolog hits in CATH, PDB, and AlphaFoldDB. Out of the 26,767 high-quality structural models, 10,493 were identified as habitat-specific, including 2,668 from aquatic marine environments, 5,290 from aquatic freshwater environments, 217 from other aquatic habitats, 22 from engineered environments, 8 from host-plants, 323 from mammals, 15 from humans, 4 from other host-associated environments, and 1,946 from terrestrial environments. Out of 47,823 medium-quality structural models, 21,460 were identified as habitat-specific. This includes 6,088 from marine environments, 10,029 from freshwater environments, 369 from other aquatic environments, 82 from engineered environments, 131 from host plants, 318 from mammals, 48 from humans, 23 from other host-associated environments, and 4,372 specific to terrestrial habitats.

For each family, the predicted fold was compared with the structural hits from these databases and combined with the gene neighborhood analysis. The F139409 family for example, which is specific to the human habitat, exhibits structural similarity to the Winged Helix DNA-binding domain (CATH: 1.10.10.10). Gene neighborhood analysis indicates that its neighboring genes predominantly encode helix-turn-helix motifs (PF01381), ERF superfamily domains (PF04404), and single-nucleotide strand DNA binding protein domains (PF00436), all of which are known for their roles in DNA binding. The helix-turn-helix domain is a well-characterized structural motif involved in DNA recognition, while the ERF superfamily includes single-strand annealing proteins. The presence of exclusively DNA-binding proteins within this gene neighborhood suggests a strong functional connection related to DNA interaction (Supplementary Figure 6A). Similarly, the F004558 family, which consists of proteins found exclusively in terrestrial environments, exhibits high structural similarity (as indicated by TM scores) to the CRAL-TRIO lipid-binding domain (CATH: 3.40.525.10). The CRAL-TRIO domain is primarily associated with lipid transport and the binding of small lipophilic molecules. Gene neighborhood analysis of the F004558 family reveals a strong correlation with membrane transport functions, with neighboring genes encoding ion channels (PF07885), sodium/calcium exchangers (PF01699), and transmembrane transport proteins (GO:0055085). Furthermore, this family is closely related to the SLOG superfamily (PF18306), which includes proteins that bind low-molecular-weight biomolecules. Notably, SLOG proteins and CRAL-TRIO domains share functional similarities as small-molecule-binding domains. This relationship extends to lipid transfer and membrane interactions, particularly in the context of membrane trafficking^24^, further reinforcing their functional link in cellular transport mechanisms (Supplementary Figure 6B).

Additionally, to focus on high-quality models, the 50 most frequent structural hits in each environment were selected, and the unique domains identified in each environment were reported (Supplementary Table 5). This analysis, along with the gene neighborhood analysis, does not pinpoint the absolutely unique structural domains for each habitat, but rather highlights the unique domains from the most frequently encountered ones. Interestingly, the most frequent and characteristic gene neighborhood functions presented in Supplementary Tables 2, 3, and 4 are closely related to the habitat-specific unique structural domains. For instance, the Carboxypeptidase-like regulatory domain (CATH: 2.60.40.1120) is found in families specific to the Terrestrial habitat, and this is further supported by neighborhood analysis, which identifies it as one of the most frequently associated functions (Pfam domain: PF13620). Another example is the STAS domain, which is strongly associated with regulation and signal transduction in bacterial systems^25^. This domain appears at the top of the list in Terrestrial-specific families (CATH: 3.30.750.24). At the same time, the gene neighborhood analysis reveals that the top Pfam domains are related to bacterial regulatory proteins (Pfam: PF00196). Considering all of the above, structural homology not only gives insights into annotating novel protein families but also underscores the significance of gene neighborhood analysis, which plays a crucial role in characterizing novel or unannotated folds.

## Discussion

The microbial world contains vast, largely unexplored protein diversity that underpins microbiome function and ecological interactions. By analyzing tens of billions of sequences from metagenomes, metatranscriptomes, and reference genomes, we substantially expand protein family space, identifying over 600,000 families with ≥100 members and 6.5 million families with ≥25 members, effectively quadrupling current coverage. Comparison of novel protein families with MAGs (Supplementary File 2) and NMPFamsDB v.1 (Supplementary File 3) reveals only limited overlap. Importantly, smaller and niche families capture rare proteins with potential functional and evolutionary significance. Our scalable framework integrates structural prediction with taxonomic, ecological, and genomic context, revealing biome-specific patterns of adaptation and uncovering both novel folds and known folds within previously uncharacterized sequence space. Beyond novel folds, we identify protein families that adopt established structural architectures but lack detectable sequence similarity to known domains.

An insightful example (protein family F431767) exemplifies this structure-guided functional inference (Figure 4). Although no similarity to Pfam domains or isolated genomes is detected, members of this family exhibit clear structural similarity to Penicillin V acylase (PVA) chain A (CATH: 3.60.60.10; PDB: 3PVA) (Figure 4A). PVA catalyzes the hydrolysis of penicillin V to 6-aminopenicillanic acid (6-APA), a key intermediate in the semi-synthesis of penicillin products in industry^26,27^. F431767 proteins are distributed primarily across marine and terrestrial environments and occur mainly within Pseudomonadota, Actinomycetota, and Chloroflexota (Figure 4C,D). The presence of this family in Chloroflexota^28^, an ecologically and metabolically versatile phylum, highlights its potential functional breadth and biotechnological relevance. Sequence comparisons with structurally characterized PVAs, guided by the penicillin V-bound structure of 2Z71, reveal conservation of catalytic features (Figure 4E), including the N-terminal nucleophile Cys12 generated by autocatalytic self-cleavage, consistent with the N-terminal nucleophile (Ntn) hydrolase superfamily^29^. Additional conserved residues (Asp30 and Asn155) contribute to substrate binding, while substitution of Tyr80 with His80 in F431767 members suggests adaptive tuning, potentially enhancing electrostatic flexibility and stability in environmentally diverse or extreme habitats. Molecular docking confirms the penicillin V model potential binding (Figure 4B).

**Figure 4.**
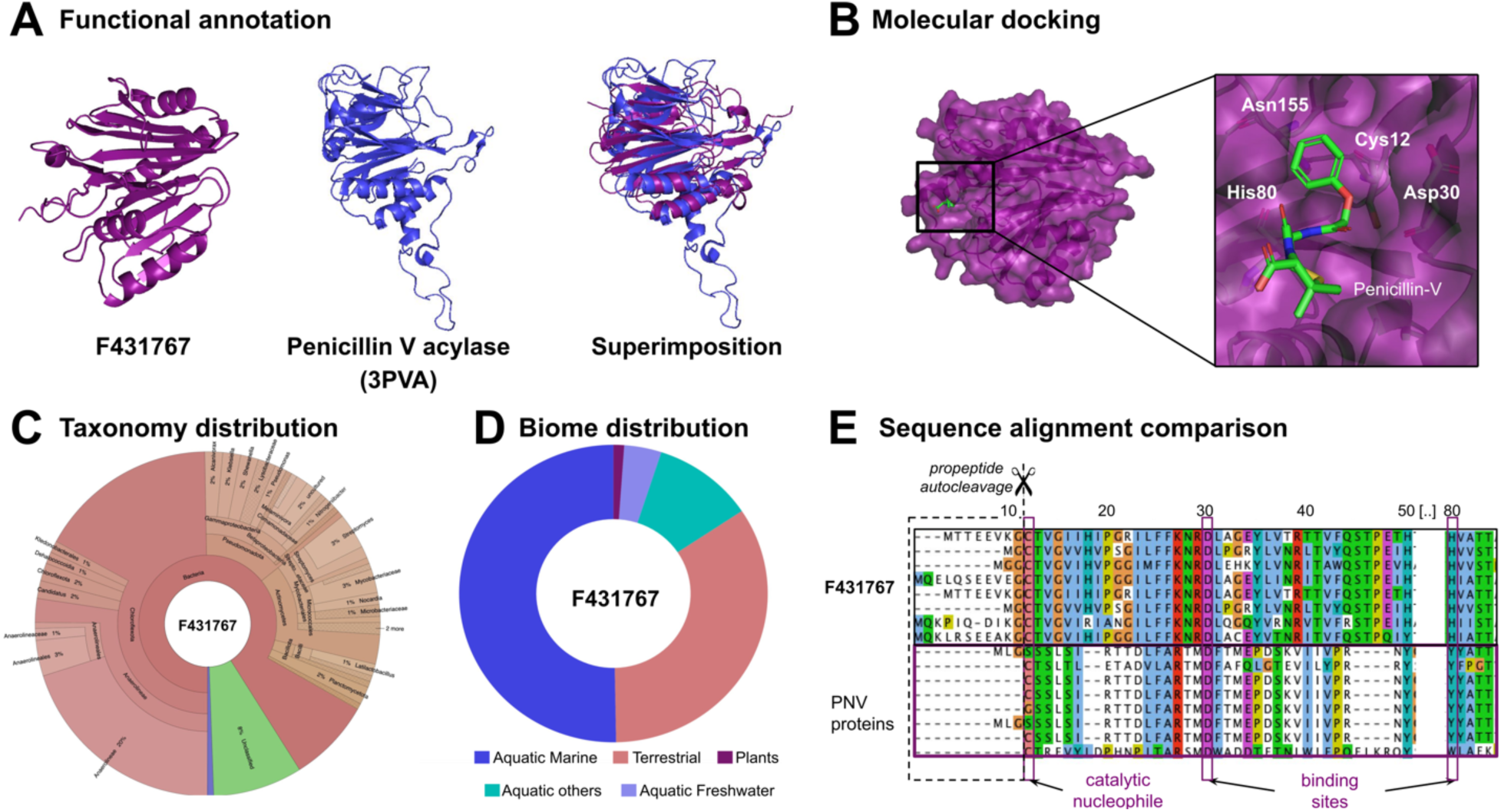
(A) Structural model of the novel protein family F431767 and its foldseek hit, penicillin V acylase (PDB code: 3PVA), including their structural superimposition. (B) Molecular docking by manual fitting of penicillin V in the F431767 structural model. Key residues conserved between penicillin V acylase and F431767 are highlighted. (C) KRONA visualization showing the taxonomic distribution of proteins belonging to the F431767 family. Members are predominantly found in Bacteria, including Pseudomonadota, Actinomycetota, and Chloroflexota. Chloroflexota includes aerobic thermophiles that utilize oxygen and thrive at high temperatures. (D) Pie chart illustrating the biome distribution of F431767 family members, with most sequences originating from aquatic marine and terrestrial environments, followed by aquatic freshwater, other aquatic environments, and plants. (E) Sequence alignment comparing penicillin V acylase PDB structures (2IWM, 2OQC, 2PVA, 2QUY, 2Z71, 3MJI, 3PVA, 4WL2) with members of the F431767 family to identify conserved and divergent residues. Key residues such as Cys12, the catalytic N-terminal nucleophile, are highly conserved, while Asp30 is also conserved and contributes to the penicillin V binding site.

Overall, this study shows that integrating global metagenomic clustering with structural and contextual annotation converts billions of uncharacterized sequences into a biome-resolved protein family landscape, providing new insights into microbial adaptation and revealing a broad reservoir of functional and biotechnological potential.

## Methods

### Data collection and filtering

All publicly accessible metagenomes, metatranscriptomes, and reference genomes were obtained from the Integrated Microbial Genomes & Microbiomes (IMG/M) database provided by the Joint Genome Institute (Figure 1). Metagenomes and metatranscriptomes were retrieved in October 2023^7^, while reference genomes were collected in December 2024. A total of 522,135,784 protein sequences were extracted from 143,797 bacterial genomes, 761,975 sequences from 19,747 viral genomes, 6,651,575 sequences from 3,128 archaeal genomes, and 9,491,122 sequences from 803 eukaryotic genomes, culminating in 539,040,456 sequences. All sequences were solely derived from isolated reference genomes, excluding metagenome-assembled genomes (MAGs) and single-amplicon genomes (SAGs). The study analyzed 49,986 microbiome datasets (40,446 metagenomes and 9,540 metatranscriptomes, totaling 10,851,187,196 scaffolds) that initially contained 40,315,976,846 proteins of varying lengths from assembled metagenomes. To maintain a dataset comprised primarily of reliable protein predictions, specific filtering criteria were employed. Given that genes situated toward the center of a contig are more likely to be complete. At the same time, those closer to the ends may be truncated; sequences located within 10 nucleotide residues of scaffold ends were also removed to mitigate the inclusion of potentially truncated genes. Furthermore, only sequences with 35 or more residues from scaffolds exceeding 500 nucleotides in length were retained to ensure that mainly complete genes were represented. This is because shorter scaffolds increase the risk that genes are located near the ends, which could lead to truncation. Low-complexity regions were masked using the tantan application (version 26)^30^. After these filtering steps, any remaining sequences shorter than 35 amino acids were also discarded. The resulting final catalog comprised 6,231,725,853 sequences derived from 44,856 microbiome datasets and 2,893,181,669 scaffolds. We consider this to be the high-quality non-redundant microbiome protein sequence set (High Quality / Non Redundant), which accounted for 15.46% of the starting dataset.

### Identification of novel proteins: hits to Pfam & Antifam

To determine which of the filtered proteins possess known domains, the filtered metagenome dataset was compared against the Pfam-A (version 37) database of profile Hidden Markov Models (HMMs) (Figure 1), which contains 21,979 families and 709 clans. Pfam hits were identified using the hmmsearch tool from the HMMER 3.3.2 package^31^, applying the default trusted cutoff. Following the removal of Pfam hits, the final dataset comprised 2,664,804,251 proteins, extracted from 44,796 datasets and 1,579,469,832 scaffolds.

The same methodology was applied for Antifam^5^, a database specifically aimed at identifying and eliminating spurious or erroneous protein sequences coming from low-complexity regions or other repetitive patterns, improperly extended open reading frames, or incorrectly translated non-protein-coding RNA genes (Figure 1). While Pfam catalogs functional protein domains and families, Antifam focuses on detecting sequences or families that are likely artifacts (such as those arising from low-complexity regions, coiled-coil structures, or other repetitive patterns) that may falsely resemble functional domains. The Antifam filtering process yielded 398,073 hits, resulting in a final dataset of 2,664,406,178 proteins (43% of the HQ-NR-MB set).

### Identification of novel proteins: hits to reference genomes

To identify novel proteins in the filtered microbiome protein dataset, additional comparisons were made against the most recent reference genomes collected in late 2024, which encompass 539,040,456 protein sequences. Two sequential methods were employed for the comparison (Figure 1). Initially, the LAST algorithm^32^ was utilized with parameters set at 30% similarity and 70% coverage^33^. LAST is designed explicitly for large-scale genomic or protein sequence comparisons, utilizing a variation of spaced seeds to enhance sensitivity while minimizing memory usage. Due to its speed and ability to handle large datasets, LAST is well-suited for metagenomic data analysis but can miss hits with lower sequence identity. For this reason, the dataset that yielded no hits in the LAST comparison (1,957,621,091 proteins) was subsequently analyzed against the reference genomes using DIAMOND ^34^, which is more sensitive, albeit slower. The DIAMOND search was performed using the same search parameters. Following these steps, the final dataset comprised 1,736,930,453 proteins, sourced from 44,721 microbiome datasets and 1,115,001,758 scaffolds.

### Clustering and generation of protein families

In this analysis, MMseqs2 Linclust^35^ was run in bidirectional mode with 30% sequence identity and 80% coverage, balancing sensitivity and computational load (Figure 1). A direct comparison of previous clustering methodology (NMPFam v.1) is presented in Supplementary File 4. Using these parameters, 370,185,087 proteins were clustered into 6,588,876 families, each containing at least 25 members. These families were derived from 44,198 datasets and spanned 253,359,325 scaffolds, representing 21.31% of the total 1,736,930,453 proteins analyzed. Proteins that did not meet the minimum clustering threshold were excluded from further analysis.

### Mapping to metagenome-assembled genomes (MAGs)

In this study, all prokaryotic metagenome-assembled genomes (MAGs) were sourced from the IMG/M database, while eukaryotic MAGs were excluded. According to the MIMAG standards^36^, the bins were classified as high, medium, or low quality, with high-quality MAGs characterized by >90% completeness and <5% contamination, as determined with CheckM^37^. As of June 2024, the dataset encompassed 21,817 high-quality prokaryotic MAGs, providing a robust foundation for the analysis of microbial diversity^38^.

### Profile generation

Novel protein families containing 100 or more members were gathered and subjected to multiple sequence alignment (MSA) generation using MAFFT^39^. After the initial alignment, trimming was applied to minimize gaps as follows: The “central” or “pivot” sequence of each MSA was identified by converting aligned sequences to numerical values, constructing an all-vs-all matrix, and computing the Hamming distance. Subsequently, the central sequence was used as a guide to identify gaps, isolate conserved regions, and ensure that poorly aligned or gapped areas were excluded from subsequent analysis. Finally, hhfilter^40^ was employed to reduce sequence redundancy within each family MSA, with parameters set to an identity threshold of 95% and a coverage of 70%, thereby reducing bias by eliminating highly similar sequences. Only families with at least 16 effective sequences after filtering were retained for structural prediction, following the protocol established in the original version of NMPFamsDB. This cutoff was chosen as it was previously found to be the minimum number of sequences per MSA capable of producing the highest percentage of high-quality results^41^.

### Candidate families for structural model prediction

Structure prediction was performed using the ColabFold^19^ implementation of AlphaFold2^42^ in *de novo* mode (no template input), following the protocol established in the original version of NMPFamsDB^41^. Five models were generated per run, with three recycle rounds computed per model. For each cluster, the model with the highest average predicted Template-Modeling (pTM) score and best average predicted Local Distance Difference Test (pLDDT) score was selected for downstream evaluation.

### Structural homology search

Only high-quality (pTM ≥ 0.7) and medium-quality (0.5 ≤ pTM < 0.7) structures were retained for further analysis. To assess structural similarity with experimentally determined structures, these models were searched using Foldseek^43^ against 5,653 CATH^21^ (v4.4, released October 2024) to detect matches to structural domains, and PDB assemblies^22^ (March 2024 release), to account for multi-domain models with matches to multimeric complexes, as well as any PDB structures not already incorporated in CATH. Hits were collected for all family structural models with an alignment TM-score greater than 0.5. For the remaining family models, query and target lengths were considered. If the query length was shorter than the target, the query alignment score was assessed with a 0.5 threshold. Conversely, for cases where the target length was shorter, a family was considered a hit if the target alignment length was at least 0.5. For the remaining families, an additional search against AlphaFoldDB^20,42^ was performed to identify models that lacked experimental structure but had been previously predicted by AlphaFold. For each hit identified in AlphaFoldDB, further annotation was carried out using metadata retrieved from the corresponding UniProt entries to determine whether the hits were functionally annotated or merely predicted.

### Superfamily generation

To assess how many families can be grouped into superfamilies based on structural information, Foldseek clustering was performed in bidirectional mode with a coverage threshold of 80% and a TM-score threshold of 0.5.

### Taxonomic annotation

The taxonomic analysis in this study focused on families containing over 100 members, adhering to criteria established in previous studies^41,44^ to ensure statistical robustness. Protein families were defined as clusters meeting this threshold, and their associated scaffolded sequences were retrieved from the IMG/M database. Taxonomic classification of these sequences began with Kraken2^45^ (version 2.1.3), a highly accurate tool for taxonomic identification. To enhance the annotation of eukaryotic proteins and minimize false positives (given the limitations of taxonomic algorithms like Kraken2 in handling eukaryotic sequences^46^), a supplementary analysis using MMseqs2 taxonomy^47^ (version 13) was performed. For scaffolds that remained unclassified after this step, the Whokaryote tool^48^ (version 1.1.2) was used to differentiate eukaryotic from prokaryotic contigs based on domain-specific gene structural features. Scaffolds still unclassified following Whokaryote analysis were further analyzed with EukRep^49^ (version 0.6.7), a tool for detecting eukaryotic sequences in metagenomic and metatranscriptomic datasets. Lastly, geNomad^50^ (version 1.8.1) was used to detect additional plasmid and viral sequences among the remaining unclassified scaffolds.

### Biome distribution

A protein family distribution across various ecosystems was performed to enhance our understanding of habitat-specific diversity within these protein families. To explore this distribution across different biomes, metadata from each environmental sample was extracted from the IMG/M database and organized using the GOLD classification system^8^.

In this study, the three primary GOLD ecosystems (*environmental, host-associated,* and *engineered*) were further divided into ten specific ecosystem types for direct comparison and enhanced detail, consistent with the NMPFamsDB study. These ten ecosystem types include *Air*, *Freshwater*, *Marine*, *Other Aquatic Environments*, *Engineered*, *Human*, *Mammal Host-associated*, *Other Host-associated Environments*, *Plants*, and *Terrestrial*. Network visualizations were created using the Gephi software^51^, with the layout generated by the Yifan-Hu algorithm^52^, while the geographic distribution was computed using each dataset’s geolocation metadata from IMG.

### Gene Neighborhood

The analysis focuses on gene families with more than 100 members. The majority of novel proteins are located close to known genes, suggesting that Pfam annotations of the known genes can provide valuable insights into the functions of the novel proteins. Specifically, 81,908,529 out of 87,393,063 scaffolds contain both novel and known genes, highlighting potential interactions or shared functions. To conduct the gene neighborhood analysis, scaffolds associated with families comprising more than 100 members were collected. These scaffolds were then examined against Pfam annotations. For each scaffold, the top five most frequent Pfam domains were identified. The Pfam domains were associated with their respective Pfam descriptions. In addition, these domains were subsequently matched to Gene Ontology (GO) terms using the InterPro2GO mapping from InterPro. To simplify and interpret the results, GO subsets (commonly referred to as GO slims), condensed versions of GO that contain curated subsets of terms, were used. The GO terms were further grouped into the three primary Gene Ontology categories: Molecular Function, Cellular Component, and Biological Process, enabling distinct, targeted analysis within each category.

## Data Availability

Both novel families from metagenomes (minimum of 25 members) and families from reference genomes and medium- to high-quality predicted structures from novel families (at least 100 members) can be found at: https://zenodo.org/records/17225887.

## Funding

Hellenic Foundation for Research and Innovation (H.F.R.I.) under the ‘Third Call for H.F.R.I. Research Projects to support faculty members and researchers’ [23592-EMISSION]; Hellenic Foundation for Research and Innovation (H.F.R.I.) under the ‘4th Call for H.F.R.I. Research Project to support Postdoctoral Researchers’ [28787-VIROMINE]; Fondation Santé; ARISE program from the European Union’s Horizon 2020 research and innovation program under the Marie Skłodowska-Curie grant agreement No 945405; Startup funds from the Penn State College of Medicine and by the Huck Innovative and Transformational Seed Fund (HITS) award from the Huck Institutes of the Life Sciences at Penn State University; Startup funds from the University of Texas at Austin; US Department of Energy Joint Genome Institute (https://ror.org/04xm1d337); US Department of Energy Office of Science user facilities, operated under contract no. DE-AC02-05CH11231; Applied Mathematics program of the DOE Office of Advanced Scientific Computing Research (DE-AC02–05CH11231).

## Notes

https://zenodo.org/records/17225887

### Competing Interest Statement

The authors have declared no competing interest.

