## Supplementary material for "Quadrupling the protein family space with global metagenomics": Supplementary File 1.pdf

**Family properties**

A detailed analysis of family properties across various family sizes (≥25, ≥50, ≥75, and ≥100 members) is offered, enabling direct comparisons between smaller and larger families (Supplementary Figure 1). Supplementary Figure 1A shows that 90% of proteins from environmental samples fall within the 35 to 200 amino acid range. In contrast, reference genomes show a greater diversity with 64% of proteins spanning a length range of 100 to 400 amino acids. Due to the fragmented nature of metagenomes, the majority of proteins are consistently shorter across all family size thresholds, whereas in reference genomes, protein lengths are more evenly distributed across larger length categories across all family size thresholds as well.

Supplementary Figure 1B illustrates that as the family size threshold increases, the number of samples or genomes contained within these families also tends to increase. While both environmental samples and reference genomes show an upward shift in sample/genome numbers with larger family sizes, reference genomes are more evenly distributed across categories with higher genome counts.

Supplementary Figure 1C shows that, as family size increases, the number of scaffolds within each family tends to grow. While most families across all size thresholds are concentrated in categories with fewer scaffolds, there is a clear shift towards higher-scaffold categories as the family size threshold increases, reflecting a similar distribution pattern observed in environmental samples. Most scaffolds are concentrated in the 1,000 to 2,500 bp range, showcasing a consistent scaffold length distribution across all categories (Supplementary Figure 1D). The scaffold length distribution reveals that scaffold lengths in metagenomes are predominantly shorter across all family size thresholds, with very few lengthy scaffolds. The minimum scaffold length accounts for 500 bp, based on the applied filtering criteria. A comparative analysis was undertaken to assess the number of scaffolds containing only novel genes versus those including known and novel genes across all family size categories. The observed pattern across different family size groups was consistent, with a notable decrease in the total number of scaffolds as family size increased. Scaffolds featuring both known and novel genes were more numerous than those containing solely novel genes. Except for scaffolds with just one gene, which constituted 5-7% of the total, most novel proteins were aligned with known proteins within the same scaffold. Scaffolds containing both known and novel genes are more prevalent, providing strong evidence for the authenticity of the novel genes identified (Supplementary Figure 1E). The presence of known genes nearby enhances opportunities for insightful gene neighborhood analyses, enriching our understanding of gene functions and interactions.

Supplementary Figure 1F shows the distribution of family sizes among families with 25 or more members. As anticipated, most families in both environmental samples and reference genomes consist of relatively few members. In detail, 68% of families from environmental samples have 25 to 50 members, compared to 46% in reference genomes. The reference genomes demonstrate a more even distribution across family sizes, reflecting the fragmented nature of metagenomic data.

Finally, the Venn diagram shown in Supplementary Figure 1G highlights that 80% of families are exclusively derived from metagenomes, reflecting their abundance, while less than 1% come solely from metatranscriptomes (100% of the members). Most notably, 20% of families are represented by both metagenomic and metatranscriptomic datasets, underscoring the considerable importance of metatranscriptomic data in capturing functional aspects of protein families. This evidence supports the assertion that these proteins are actively translated, demonstrating their relevance across all family-size categories.

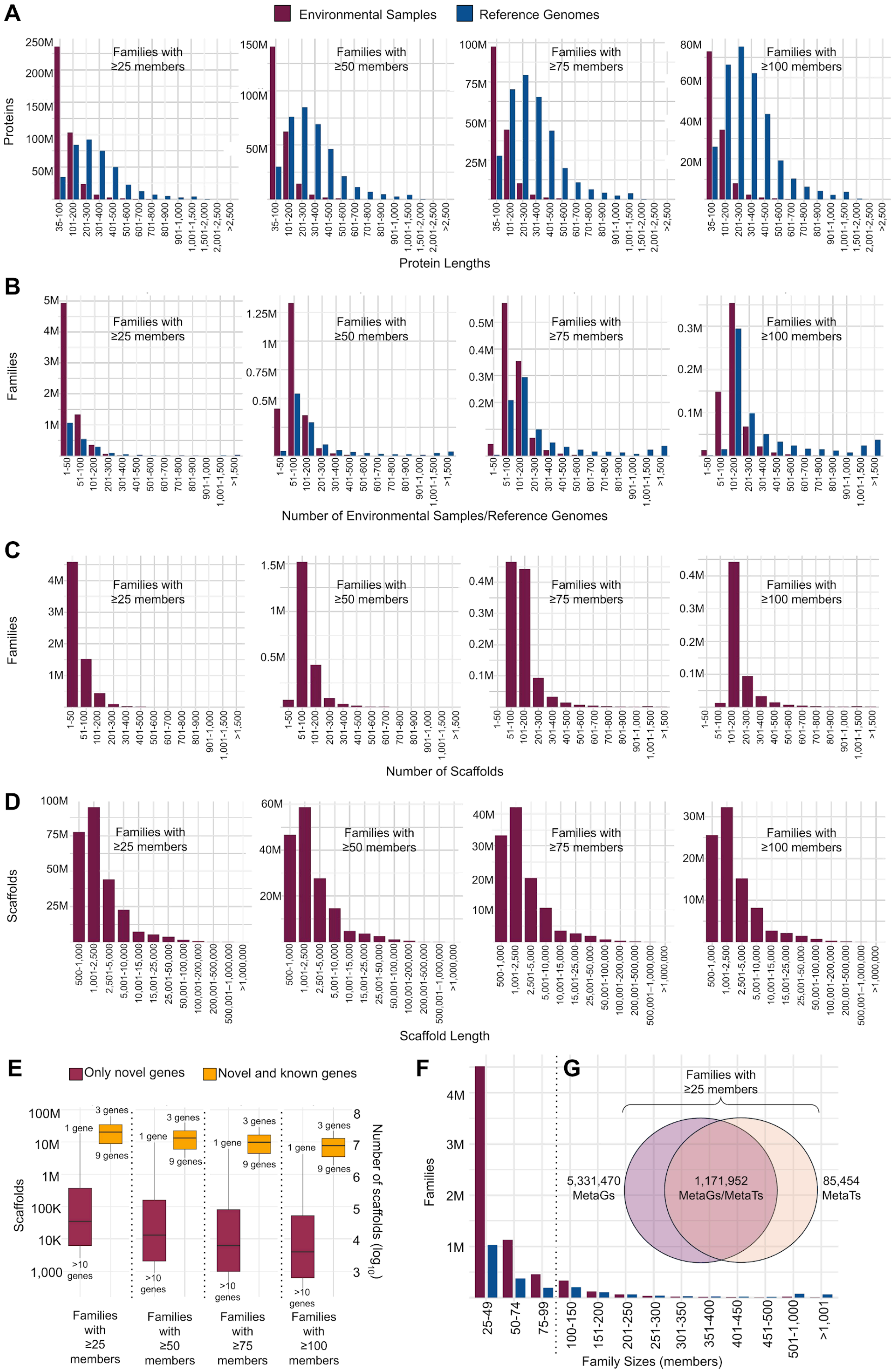

A
Environmental Samples
Reference Genomes
Proteins
Families with ≥25 members
Families with ≥50 members
Families with ≥75 members
Families with ≥100 members
250M
200M
150M
100M
50M
75M
25M
80M
60M
40M
20M
35-100
101-200
201-300
301-400
401-500
501-600
601-700
701-800
801-900
901-1,000
1,001-1,500
1,501-2,000
2,001-2,500
>2,500
Protein Lengths
B
Families
5M
4M
3M
2M
1M
1.25M
0.75M
0.5M
0.4M
0.3M
0.2M
0.1M
1-50
51-100
101-200
201-300
301-400
401-500
501-600
601-700
701-800
801-900
901-1,000
1,001-1,500
>1,500
Number of Environmental Samples/Reference Genomes
C
Families
1.5M
Number of Scaffolds
D
Scaffolds
100M
75M
50M
25M
60M
40M
20M
30M
10M
500-1,000
1,001-2,500
2,501-5,000
5,001-10,000
10,001-15,000
15,001-25,000
25,001-50,000
50,001-100,000
100,001-200,000
200,001-500,000
500,001–1,000,000
>1,000,000
Scaffold Length
E
Only novel genes
Novel and known genes
Scaffolds
100M
10M
1M
100K
10K
1,000
1 gene
3 genes
9 genes
>10 genes
Number of scaffolds (log10)
8
7
6
5
4
3
Families with ≥25 members
Families with ≥50 members
Families with ≥75 members
Families with ≥100 members
F
Families
4M
3M
2M
1M
25-49
50-74
75-99
100-150
151-200
201-250
251-300
301-350
351-400
401-450
451-500
501-1,000
>1,001
Family Sizes (members)
G
Families with ≥25 members
5,331,470 MetaGs
1,171,952 MetaGs/MetaTs
85,454 MetaTs

**Supplementary Figure 1.** Family properties for various sizes (≥25, ≥50, ≥75, and ≥100 members). **(A)** Distribution of protein lengths for environmental sample proteins compared to reference genome proteins. **(B)** Distribution of family sample counts for environmental samples and reference genomes. **(C)** Number of scaffolds per environmental sample family. **(D)** Distribution of scaffold lengths within environmental sample families. **(E)** Comparison of scaffolds containing only novel genes versus scaffolds containing both novel and known genes. Scaffolds are categorized by gene count, ranging from single-gene scaffolds to those with 10 or more genes. **(F)** Family size distribution for families with ≥25 members across environmental samples and reference genomes. **(G)** Venn diagram for the families with at least 25 members showing the overlap of family compositions derived solely (100% of the members) from metagenomes (MetaGs), metatranscriptomes (MetaTs), or both.
