## Supplementary material for "Quadrupling the protein family space with global metagenomics": Supplementary File 2.pdf

### Comparison with the MAG catalog

To investigate novelty further, we compared the newly identified protein families against high-quality Metagenome Assembled Genome (MAG) catalogs. We collected 21,817 high-quality MAGs from IMG/M and utilized DIAMOND for comparison with the representative sequences of novel protein families, applying criteria of 30% identity and 70% coverage (Table 2).

**Supplementary Table 6.** Comparison of novel protein families generated from metagenomic and metatranscriptomic data at varying family sizes (≥25, ≥50, ≥75, and ≥100 members) with MAGs.

| **Environmental Samples** | | | | |
|---|---|---|---|---|
| Proteins for clustering | 1,753,296,308 | | | |
| Family size | ≥25 members | ≥50 members | ≥75 members | ≥100 members |
| Families | 6,588,876 | 2,160,632 | 1,052,590 | 608,258 |
| Proteins | 370,185,087 | 221,646,880 | 155,259,616 | 117,330,466 |
| **Comparison with MAGs** | | | | |
| High-Quality MAGS | 21,817 | | | |
| Using ***DIAMOND*** (30% identity & 70% coverage bidirectionally) | | | | |
| Hits to MAGs | 1,075,130 | 443,338 | 247,314 | 157,471 |
| % hits MAGs | 16.32% | 20.52% | 23.50% | 25.89% |
| %MAGs Coverage | 100% | 98.35% | 94.56% | 89.35% |

For families with at least 25 members, 1,075,130 of the 6,588,876 representative sequences (16.32%) matched entries in MAGs catalogs, corresponding to 21,816 out of 21,817 MAGs, resulting in 100% coverage. As the family size threshold increased, resulting in fewer families, the representation of MAGs in families improved. In families with at least 50 members, around 20.5% of novel families matched MAGs, covering about 98.35% of the database. For families with 75 or more members, approximately 23.5% of novel families aligned with MAGs, covering roughly 94.56% of the MAGs catalog. In families with 100 or more members, about 25.89% of novel families matched MAGs, covering about 89.35% of them.

After excluding the MAG hits, the remaining novel protein space was determined to include 1,702,721,971 proteins. While the proteins from the MAGs maintain their novelty, additional clustering after removing the proteins that have hits to MAGs highlights minimal changes - less than 10% in the total number of families and proteins (Supplementary Table 7).

**Supplementary Table 7:** Comparison of novel protein clusters generated from metagenomic and metatranscriptomic data after removing MAGs hits at varying cluster sizes (≥25, ≥50, ≥75, and ≥100 members) with isolates genomes and NMPFamsDB (version 1) using two different methods.

| **Reference Genomes** | | | | |
|---|---|---|---|---|
| Proteins for clustering [illegible] | | | | |
| Clustering size | ≥25 members | ≥50 members | ≥75 members | ≥100 members |
| Clusters | 2,229,233 | 1,200,486 | 824,621 | 630,433 |

| | | | | |
|---|---|---|---|---|
| Number of Genomes | 155,857 | 154,072 | 153,253 | 152,506 |
| Proteins | 393,429,871 | 358,091,618 | 335,371,741 | 318,713,491 |
| % of proteins | 72.99% | 66.43% | 62.22% | 59.13% |
| **Environmental Samples** | | | | |
| Proteins for clustering | | | | |
| Clustering size | ≥25 members | ≥50 members | ≥75 members | ≥100 members |
| Clusters | 5,867,110 | 1,778,233 | 867,075 | 499,600 |
| Number of Metagenomes | 44,030 | 43,371 | 42,853 | 42,391 |
| Scaffolds | 212,490,117 | 129,720,097 | 93,476,007 | 71,921,521 |
| Proteins | 311,647,967 | 182,731,897 | 128,095,924 | 96,737,284 |
| % of proteins | 18.75% | 10.99% | 7.70% | 5.82% |
| Comparison with reference genomes | | | | |
| **% of cluster increase** | **263%** | **148%** | **105%** | **79%** |
| % of protein increase | 79% | 51% | 38% | 30% |

| **Comparison with NMPFamsDB (v1.0)** | | | | |
|---|---|---|---|---|
| **NMPFamsDB (version 1)** | **106,198** | | | |
| Method1: Using ***DIAMOND*** (30% identity & 70% coverage bidirectionally) | | | | |
| Hits to NMPFamsDB | 525,918 | 248,027 | 165,325 | 120,193 |
| % hits NMPFamsDB | 8.96% | 13.95% | 19.07% | 24.06% |
| %NMPFamsDB Coverage | 69.27% | 57.62% | 51.18% | 45.27% |
| Method 2: Using ***HMM*** search | | | | |
| Hits to NMPFamsDB | 928,396 | 396,607 | 248,503 | 172,898 |
| % hits NMPFamsDB | 15.82% | 22.30% | 28.66% | 34.61% |
| %NMPFamsDB Coverage | 89.74% | 79.30% | 72.92% | 67.39% |
