## Supplementary material for "Quadrupling the protein family space with global metagenomics": Supplementary File 3.pdf

**Comparison with NMPFamsDB v.1**

This study introduces an advanced approach to uncovering novel protein families (or "dark matter") from a diverse set of globally collected metagenomes and metatranscriptomes, specifically targeting proteins without matches to any known reference genome or Pfam entry. Novel families were reported across various sizes (≥25, ≥50, ≥75, and ≥100 members), although the primary focus was on families with 100 or more members to align with the previous NMPFamsDB study and to ensure statistical significance. This ≥100-member threshold minimizes the likelihood of random clustering, especially given the extensive global diversity of the samples.

In contrast to the NMPFamsDB (version 1.0)[16], which relied on sequence similarity networks and graph-based HipMCL clustering, this study utilized MMseqs2, a heuristic-based tool that offers faster processing and scalability for large datasets. By reducing the member similarity threshold within a family from 70% (used in the NMPFamsDB v1.0) to 30%, the study improves the structural sensitivity and comprehensiveness of the clustering scheme, and increases the diversity among the different reported novel families. Although every novel protein family in NMPFamsDB (version 1.0) displays an average in-family identity of at least 70%, the updated dataset for families with 100 or more members now includes a 5% increase in clusters with an average identity below 70%. This update covers 33,525 families whose average identities range between 30% and 70%.

Additionally, including smaller families with a minimum of 25 members expanded the analysis to 6.5 million families, a substantial increase over the 608,258 families of ≥100 members or the 106,000 families stored in the NMPFamsDB catalog. This refined methodology significantly increases the proportion of proteins within the dataset that are assigned to families, improving coverage from 13.66% in NMPFamsDB to 21.52%.

The comparison of representative sequences of the newly found novel families against clusters stored in NMPFamsDB was performed using DIAMOND, with criteria of 30% identity and 70% coverage (bidirectionally), and an HMM search applying trusted cutoffs (Table 3). For clusters with at least 25 members, 321,087 of the 6,588,876 representative sequences (4.78%) matched entries in NMPFamsDB, corresponding to 74,286 out of 106,198 families, resulting in 69.95% coverage. The HMM search yielded even better outcomes, owing to its ability to detect remote homologs, reaching 84.66% coverage of the database families. Additionally, using the HMM search, 6.35% of novel protein families from this study have hits to NMPFamsDB. As the clustering threshold increased, resulting in fewer clusters, the representation of NMPFamsDB families in clusters improved. In clusters with at least 50 members, around 10% of novel families matched NMPFamsDB families, covering about 70% of the database. For clusters with 75 or more members, approximately 12% of novel families aligned with NMPFamsDB, covering roughly 65% of the database. In clusters with 100 or more members, about 15% of novel families matched NMPFamsDB, covering about 60% of its families.

An interesting observation emerges on the overlap of families between the old and new datasets, as both studies explored environmental samples from IMG/M. Roughly 85% of NMPFamsDB families are included in the new dataset, with the remaining 15% either clustered into smaller groups or left out, likely due to their recognition as Pfam domains or reference genomes. Although 60% of the NMPFamsDB families are found in clusters with more than 100 members, 25% are spread across smaller clusters. These outcomes align with expectations, given the varying clustering techniques and the significant growth of the dataset. Notably, this overlap does not reduce the novelty of the new families identified, as only 15% overlap with known families, highlighting the considerable expansion of the dataset and its role in uncovering novel protein families. A comparison between NMPFamsDB and the latest Pfam release (v37.0) was conducted using an HMM search with trusted cutoffs. This analysis identified 8,246 NMPFams matching 2,554 Pfam profiles, confirming that NMPFams represent authentic protein domains. A comparison of NMPFams with MAGs further supports this fact. Using DIAMOND (30% identity and 70% coverage) and HMM search with trusted cutoffs, it was found that approximately half of the 106,198 NMPFams (~50%) have matches in the MAGs catalog.

**Supplementary Table 8.** Comparison of novel protein clusters generated from metagenomic and metatranscriptomic data at varying cluster sizes (≥25, ≥50, ≥75, and ≥100 members) with NMPFamsDB (version 1) using two different methods.

| | | | | |
|---|---|---|---|---|
| **Environmental Samples** | | | | |
| Proteins for clustering | 1,753,296,308 | | | |
| Clustering size | ≥25 members | ≥50 members | ≥75 members | ≥100 members |
| Clusters | 6,588,876 | 2,160,632 | 1,052,590 | 608,258 |
| Proteins | 370,185,087 | 221,646,880 | 155,259,616 | 117,330,466 |
| **Comparison with NMPFamsDB (v1.0)** | | | | |
| NMPFamsDB (version 1) | 106,198 | | | |
| Method1: Using ***DIAMOND*** (30% identity & 70% coverage bidirectionally) | | | | |
| Hits to NMPFamsDB | 321,087 | 169,638 | 110,909 | 80,183 |
| % hits NMPFamsDB | 4.87% | 7.85% | 10.54% | 13.18% |
| %NMPFamsDB Coverage | 69.95% | 56.89% | 47.89% | 40.80% |
| Method 2: Using ***HMM*** search | | | | |
| Hits to NMPFamsDB | 418,106 | 209,528 | 132,509 | 93,460 |
| % hits NMPFamsDB | 6.35% | 9.70% | 12.59% | 15.37% |
| %NMPFamsDB Coverage | 84.66% | 74.08% | 66.35% | 60.11% |
