## Supplementary material for "Quadrupling the protein family space with global metagenomics": Supplementary File 4.pdf

**MMseqs Linclust over other MMseqs clustering modes**

In our previous study[1,2], we generated clusters for both reference genomes and novel clusters hosted in the NMPFamsDB database by first constructing an all-versus-all similarity matrix and then keeping every pairwise alignment with >70% identity over 80% coverage bidirectionally. Subsequently, we applied the HipMCL graph-based clustering algorithm[3] to the resulting sequence similarity network (SSN), utilizing high-performance computational resources. In the current study, we explored the clustering capabilities offered by MMseqs2, selected for its scalability and linear runtime complexity, to evaluate which algorithms provide results comparable to those used for NMPFamsDB (SSN + HipMCL). MMseqs2 mainly presents two clustering approaches: *(i)* Linclust and *(ii)* cascading clustering mode, each with inherent advantages and disadvantages. Linclust operates rapidly by utilizing a k-mer strategy, making it particularly effective for efficiently clustering large-scale datasets, though this speed may come at a cost to sensitivity. In contrast, the cascading clustering mode relies on traditional exhaustive methods, performing comprehensive pairwise alignments that enhance sensitivity and suitability for identifying low-similarity clusters, albeit at a slower processing speed which may limit its application in high-throughput scenarios. Both methods in MMseqs2 can be employed in two specific configurations: *(i)* Greedy Set Cover and *(ii)* Connected Component, thereby enabling optimization based on desired cluster granularity and network connectivity requirements. Benchmarking these methods against HipMCL illustrates that MMseqs2 offers scalability and speed and maintains sensitivity, establishing it as a viable alternative to HipMCL for large-scale biological clustering endeavors.

To evaluate the similarity of MMseqs2 to HipMCL in clustering protein families, we benchmarked both methods using reference genomes previously clustered for NMPFamsDB. This comparison aimed to replicate HipMCL's clustering patterns, particularly focusing on forming protein families with over 100 members, as in the original clustering, to maintain consistency in family sizes and grouping trends across methods. When using MMseqs2's Linclust in Greedy Set Cover mode, 78.62% of the dataset clustered into groups exceeding 100 members, a proportion that increased to 90.64% with Connected Component mode. Similarly, the MMseqs2 Cluster algorithm in Greedy Set Cover mode grouped approximately 80% of proteins into families with over 100 members, rising to 93% under Connected Component mode. Notably, Linclust produced clustering results comparable to those of the Cluster algorithm but completed the task in a third of the time, making it the preferred choice for its substantial efficiency benefits.

To identify the most effective Linclust mode for our dataset, we evaluated both algorithms on reference genomes and environmental samples. For the reference genomes, Linclust's Greedy Set Cover mode clustered 59% of the total proteins into 630,433 families, each containing over 100 members. In comparison, the Linclust Connected Component mode formed 307,852 families, covering 75% of the dataset. For metagenomic samples, the Greedy Set Cover mode produced 608,258 families, accounting for 6.76% of the proteins, whereas the Connected Component mode generated 891,625 families, encompassing 15.77% of the protein dataset.

In the Linclust Greedy Set Cover mode, the largest cluster contains 10,027 members, with only 0.08% of clusters holding more than 2,000 members. Conversely, the Linclust Connected Components mode produced the largest cluster with 12,641 members, whereas approximately 1.4% contained more than 2,000 members. Given that both methods perform similarly in HipMCL comparison parameters and processing speed, we opted for the Linclust Greedy Set Cover mode.
