## Supplementary material for "Quadrupling the protein family space with global metagenomics": Supplementary Table 1.pdf

| Database | Type / Focus | Key Features | Number of Entries (as of March 2025) |
|---|---|---|---|
| **UniProtKB** [1] | Protein sequences & functional annotations | Two components: 1) Swiss-Prot (reviewed, manually curated), 2) TrEMBL (unreviewed, automated) | 253 million protein sequences |
| **Protein Data Bank (PDB)** [2] | 3D macromolecular structures | Experimental structural data for proteins and complexes | 232,418 protein structures |
| **RefSeq** [3] | Genomes, transcripts, proteins | Curated reference sequences across multiple organisms | 391,903,900 proteins from 162,138 organisms |
| **GenBank** [4] | Nucleotide sequences (includes proteins) | Publicly available sequences from diverse organisms | 4,7 billion sequences |
| **IMG/M v7.0** [5] | Microbial genomics & metagenomics | Supports analysis of microbial genomes, metagenomes, and metatranscriptomes | 544 million genes (genomes) & 82 billion genes (assembled/unassembled metagenomes & metatranscriptomes) |
| **UniParc** [1,6] | Non-redundant protein sequences | Comprehensive archive integrating multiple sources | 916 million protein sequences |
| **Big Fantastic Database (BFD)** [7] | Protein sequence database | Large-scale protein sequence repository (Uniprot/TrEMBL+Swissprot, Metaclust, and Soil Reference Catalog Marine Eukaryotic Reference Catalog assembled by Plass) | 2.5 billion protein sequences clustered |
| **AlphaFoldDB** [8] | Predicted protein structures | Computationally predicted 3D structures | over 200 million entries |
| **MGnify** [9] | Metagenomic protein sequences | Focuses on protein sequences derived from metagenomic studies | 24 billion protein sequences |
| **ColabFold** [10] | Extends BFD & MGnify | Representative protein sequences & total members | 209,335,865 representative sequences & 738,695,580 total members |
| **InterPro** [11] | Protein family & functional analysis | Integrates multiple databases including Pfam, SMART | Over 200 million protein sequences annotated |

| | | | |
|---|---|---|---|
| **COGs** [12] | Clusters of Orthologous Groups | Groups orthologous proteins from Bacteria and Archaea: evolutionary classification | 5,640,669 protein sequences in 5,050 COG clusters |
| **KOG** [13] | Eukaryotic orthologous groups | Groups orthologous proteins from eukaryotes (animals, fungi, plants, and microsporidia): evolutionary classification | 59,838 protein sequences in 4,852 KOG clusters |
| **NMPFamsDB** [14,15] | Novel metagenome protein families with no hits to Pfam and reference proteomes | Derived from metagenomic/metatranscriptomic data | 106,000 protein families: (20 million proteins) with ≥100 members each |
| **MetaVR** [16] | A large collection of Uncultivated Viral Genomes (UViGs) and viral protein clusters from microbiomes and isolate genomes | Derived from metagenomes, metatranscriptomes, isolates, SAGs and MAGs | 383,034 viral protein clusters with more than 100 members, 748,907 3D models (42,390,306 protein sequence) |
