## Supplementary material for "Quadrupling the protein family space with global metagenomics": Supplementary Table 2.pdf

**Supplementary Table 2:** Biome-specific Pfam descriptions, including the corresponding Pfam ID and the number of families in which each Pfam appears.

| NAME | GO ID | Family count |
|---|---|---|
| **Humans** | | |
| MatE | PF01554 | 616 |
| LytTr DNA-binding domain | PF04397 | 199 |
| Peptidase family M23 | PF01551 | 181 |
| AraC-like ligand binding domain | PF02311 | 161 |
| Probable transposase | PF01385 | 158 |
| RNA pseudouridylate synthase | PF00849 | 154 |
| Phage antirepressor protein KilAC domain | PF03374 | 145 |
| Family of unknown function(DUF5977) | PF19404 | 132 |
| Putative binding domain,N-terminal | PF13004 | 128 |
| MFS/sugar transport protein | PF13347 | 122 |
| Fibronectin type III-like domain | PF14310 | 104 |
| Rnf-Nqr_subunit,_membrane_protein | PF02508 | 98 |
| **Engineered** | | |
| Secretion system C-terminal sorting domain | PF18962 | 114 |
| 4Fe-4S binding domain | PF12837 | 114 |
| Metallo-beta-lactamase superfamily | PF14597 | 106 |
| FlgD Ig-like domain | PF13860 | 72 |
| Tripartite ATP-independent periplasmic transporter,DctM component | PF06808 | 69 |
| Prokaryotic N-terminal methylation motif | PF07963 | 64 |
| Bacterial extracellular solute-binding protein, family 7 | PF03480 | 55 |
| Phosphoribosyl transferase domain | PF00156 | 52 |
| EamA-like transporter family | PF00892 | 44 |
| Thioredoxin-like [2Fe-2S] ferredoxin | PF01257 | 40 |
| OmpA family | PF00691 | 40 |
| HAMP domain | PF18947 | 33 |
| **Terrestrial** | | |
| Carboxypeptidase regulatory-like domain | PF13620 | 3516 |
| Bacterial regulatory proteins, luxR family | PF00196 | 2800 |
| Glyoxalase/Bleomycin resistance protein/Dioxygenase superfamily | PF13669 | 2678 |
| Tripartite tricarboxylate transporter family receptor | PF03401 | 2553 |
| KR domain | PF08659 | 1962 |

| | | |
|---|---|---|
| PilZ domain | PF07238 | 1718 |
| Alcohol dehydrogenase GroES-like domain | PF08240 | 1649 |
| Aldo/keto reductase family | PF00248 | 1531 |
| Cytochrome c | PF00034 | 1327 |
| NUDIX domain | PF14815 | 1252 |
| Acyl-CoA dehydrogenase, C-terminal domain | PF00293 | 1252 |
| Acyl-CoA dehydrogenase, C-terminal domain | PF21263 | 1099 |
| **Aquatic Others** | | |
| Putative restriction endonuclease | PF05685 | 451 |
| Protein of unknown function (DUF1559) | PF07596 | 368 |
| Ribbon-helix-helix protein, copG family | PF01402 | 228 |
| ParB/Sulfiredoxin domain | PF02195 | 191 |
| Helicase HerA, central domain | PF01935 | 138 |
| AAA ATPase domain | PF13191 | 133 |
| Nucleotidyltransferase domain | PF01909 | 131 |
| Transposase DDE domain | PF13751 | 126 |
| PEP-CTERM motif | PF07589 | 122 |
| Putative transposase DNA-binding domain | PF07282 | 99 |
| **Aquatic Marine** | | |
| Putative 2OG-Fe(II) oxygenase | PF13759 | 4514 |
| PDDEXK-like domain of unknown function (DUF3799) | PF12684 | 3347 |
| DnaB-like helicase C terminal domain | PF03796 | 3190 |
| Terminase small subunit | PF03592 | 3156 |
| Cell Wall Hydrolase | PF07486 | 2996 |
| Staphylococcal nuclease homologue | PF00565 | 2698 |
| Thymidylate synthase complementing protein | PF02511 | 2427 |
| Glutaredoxin | PF00462 | 2332 |
| 5'-3' exonuclease, N-terminal resolvase-like domain | PF02739 | 2227 |
| Hsp20/alpha crystallin family | PF00011 | 2033 |
| Holin of 3TMs, for gene-transfer release | PF11351 | 1968 |
| Ribonucleotide reductase, small chain | PF00268 | 1647 |
| **Aquatic Freshwater** | | |
| Transcription factor WhiB | PF02467 | 4693 |
| Phage portal protein | PF04860 | 4012 |

| | | |
|---|---|---|
| Endodeoxyribonuclease RusA | PF05866 | 3327 |
| Caudovirus prohead serine protease | PF04586 | 3247 |
| Concanavalin A-like lectin/glucanases superfamily | PF13385 | 3241 |
| YqaJ-like viral recombinase domain | PF09588 | 3163 |
| Transglycosylase SLT domain | PF01464 | 2981 |
| D-alanyl-D-alanine carboxypeptidase | PF13539 | 2642 |
| Ferredoxin-like domain in Api92-like protein | PF18406 | 2337 |
| Family of unknown function (DUF5856) | PF19174 | 2231 |
| Methyltransferase FkbM domain | PF05050 | 2162 |
| Chitinase class I | PF00182 | 1957 |
| Domain of unknown function (DUF6378) | PF19905 | 1936 |
| Phage capsid family | PF05065 | 1929 |
| **Mammals** | | |
| Family of unknown function (DUF5662) | PF18907 | 122 |
| GIY-YIG catalytic domain | PF20815 | 120 |
| GIY-YIG catalytic domain | PF01541 | 120 |
| BspA type Leucine rich repeat region (6 copies) | PF13306 | 105 |
| ADP-ribosylation factor family | PF00025 | 91 |
| 4Fe-4S single cluster domain | PF13459 | 77 |
| ATP dependent DNA ligase domain | PF01068 | 71 |
| dUTPase | PF08761 | 66 |
| SprT-like family | PF10263 | 60 |
| Antidote-toxin recognition MazE, bacterial antitoxin | PF04014 | 54 |
| Resolvase, N terminal domain | PF00239 | 53 |
| Terminase large subunit, T4likevirus-type, N-terminal | PF03237 | 43 |
| ATP cone domain | PF03477 | 43 |
| YodL-like | PF14191 | 41 |
| Tetrahydrofolate dehydrogenase/cyclohydrolase, catalytic domain | PF00763 | 36 |
| RNase H | PF00075 | 35 |
| Phage gp6-like head-tail connector protein | PF05135 | 28 |
| Polypeptide deformylase | PF01327 | 27 |
| **Plants** | | |
| Reverse transcriptase-like | PF13456 | 1124 |
| gag-polypeptide of LTR copia-type | PF14244 | 1076 |
| Putative gypsy type transposon | PF04195 | 866 |

| | | |
|---|---|---|
| Transposase family tnp2 | PF02992 | 666 |
| GAG-pre-integrase domain | PF13976 | 630 |
| Domain of unknown function (DUF4218) | PF13960 | 565 |
| Plant mobile domain | PF10536 | 414 |
| PPR repeat | PF12854 | 392 |
| Domain of unknown function (DUF4216) | PF13952 | 335 |
| Proton-conducting membrane transporter | PF00361 | 246 |
| Domain of unknown function (DUF4283) | PF14111 | 243 |
| NB-ARC domain | PF00931 | 230 |
| NADH dehydrogenase | PF00146 | 182 |
| ABC-2 type transporter | PF19055 | 181 |
| Mitovirus RNA-dependent RNA polymerase | PF05919 | 174 |
| Possible lysine decarboxylase | PF03641 | 164 |
| Kinesin motor domain | PF00225 | 157 |
| Protein phosphatase 2C | PF13672 | 136 |
| Ribosomal protein S7p/S5e | PF00177 | 133 |
| GDSL-like Lipase/Acylhydrolase | PF16255 | 119 |
| No apical meristem (NAM) protein | PF02365 | 102 |
| Cytochrome C assembly protein | PF01578 | 91 |
| **Host others** | | |
| Chromo (CHRromatin Organisation MOdifier) domain | PF00385 | 217 |
| Aspartyl protease | PF13650 | 210 |
| Aspartyl protease | PF09668 | 210 |
| Zinc knuckle | PF15288 | 71 |
| Endonuclease-reverse transcriptase | PF14529 | 47 |
| Transposase (partial DDE domain) | PF01359 | 40 |
| PIF1-like helicase | PF05970 | 32 |
| HTH domain in Mos1 transposase | PF17906 | 32 |
| NACHT domain | PF05729 | 29 |
| Ty3 transposon capsid-like protein | PF19259 | 26 |
| GMC oxidoreductase | PF05199 | 23 |
| Helitron helicase-like domain at N-terminus | PF14214 | 22 |
| Chitin synthase | PF03142 | 22 |
| Helix-turn-helix domain (DUF4817) | PF16087 | 21 |
| Aldehyde dehydrogenase family | PF00171 | 18 |

| | | |
|---|---|---|
| short chain dehydrogenase | PF00106 | 17 |
| Hsp70 protein | PF00012 | 15 |
| 50S ribosome-binding GTPase | PF01926 | 15 |
| Ferrous iron transport protein B | PF02421 | 14 |
| DnaJ domain | PF00226 | 12 |
| ROK family | PF00480 | 11 |
