## Supplementary material for "Quadrupling the protein family space with global metagenomics": Supplementary Table 3 .pdf

**Supplementary Table 3:** Biome-specific biological processes, including the corresponding Gene Ontology ID and the number of families in which each Gene Ontology ID appears.

| NAME | GO ID | Family count |
|---|---|---|
| **Mammals** | | |
| tetrahydrofolate biosynthetic process | GO:0046654 | 20 |
| tRNA modification | GO:0006400 | 20 |
| translational termination | GO:0006415 | 20 |
| sporulation resulting in formation of a cellular spore | GO:0030435 | 14 |
| obsolete organic substance metabolic process | GO:0071704 | 14 |
| protein modification process | GO:0036211 | 10 |
| queuosine salvage | GO:1990397 | 9 |
| sodium-dependent phosphate transport | GO:0044341 | 8 |
| negative regulation of DNA -templated transcription | GO:0045892 | 7 |
| protein homooligomerization | GO:0051260 | 7 |
| microtubule-based process | GO:0007017 | 7 |
| regulation of cell shape | GO:0008360 | 6 |
| **Terrestrial** | | |
| DNA damage response | GO:0006974 | 547 |
| metal ion transport | GO:0030001 | 546 |
| protein-DNA covalent cross-linking repair | GO:0106300 | 538 |
| riboflavin biosynthetic process | GO:0009231 | 491 |
| fatty acid metabolic process | GO:0006631 | 408 |
| signal peptide processing | GO:0006465 | 307 |
| asparagine biosynthetic process | GO:0006529 | 284 |
| protein transport | GO:0015031 | 274 |
| **Aquatic Freshwater** | | |
| chitin catabolic process | GO:0006032 | 1957 |
| DNA metabolic process | GO:0006259 | 1111 |
| spermidine biosynthetic process | GO:0008295 | 1087 |
| virion assembly | GO:0019068 | 1003 |
| superoxide metabolic process | GO:0006801 | 537 |
| response to oxidative stress | GO:0006979 | 362 |
| peptidoglycan turnover | GO:0009254 | 268 |
| **Aquatic Others** | | |
| DNA-mediated transformation | GO:0009294 | 42 |

| | | |
|---|---|---|
| photosynthesis, light reaction | GO:0019684 | 33 |
| DNA replication, synthesis of primer | GO:0006269 | 31 |
| methylation | GO:0032259 | 30 |
| archaeal or bacterial-type_flagellum-dependent cell motility | GO:0097588 | 19 |
| photosynthetic electron transport in photosystem II | GO:0009772 | 14 |
| Mo-molybdopterin cofactor biosynthetic process | GO:0006777 | 11 |
| **Humans** | | |
| beta-lactam antibiotic catabolic process | GO:0030655 | 52 |
| antibiotic catabolic process | GO:0017001 | 52 |
| quorum sensing | GO:0009372 | 46 |
| purine nucleotide biosynthetic process | GO:0006164 | 44 |
| L-lysine transmembrane transport | GO:1903401 | 27 |
| **Aquatic Marine** | | |
| chromosome organization | GO:0051276 | 3181 |
| viral process | GO:0016032 | 1363 |
| protein glycosylation | GO:0006486 | 569 |
| nucleobase-containing compound metabolic process | GO:0006139 | 532 |
| **Engineered** | | |
| isoprenoid biosynthetic process | GO:1902768 | 17 |
