## Supplementary material for "Quadrupling the protein family space with global metagenomics": Supplementary Table 4 .pdf

**Supplementary Table 4:** Biome-specific molecular functions, including the corresponding Gene Ontology ID and the number of families in which each Gene Ontology ID appears.

| NAME | GO ID | Family count |
|---|---|---|
| **Host others** | | |
| aspartic-type endopeptidase activity | GO:0004190 | 17 |
| ubiquitin-protein transferase activity | GO:0004842 | 13 |
| translation elongation factor activity | GO:0003746 | 9 |
| protein-N(PI)-phosphohistidine-sugar phosphotransferase activity | GO:0008982 | 8 |
| transferase activity | GO:0016740 | 7 |
| rRNA binding | GO:0019843 | 6 |
| protein-folding chaperone binding | GO:0051087 | 6 |
| mannosyl-oligosaccharide 1,2-alpha mannosidase activity | GO:0004571 | 6 |
| alanine-tRNA ligase activity | GO:0004813 | 6 |
| 1-deoxy-D-xylulose-5-phosphate synthase activity | GO:0008661 | 6 |
| hydro-lyase activity | GO:0016836 | 5 |
| **Humans** | | |
| chromate transmembrane transporter activity | GO:0015109 | 83 |
| **Terrestrial** | | |
| cyclic-di-GMP binding | GO:0035438 | 1718 |
| transaminase activity | GO:0008483 | 602 |
| **Aquatic Others** | | |
| nucleotidyltransferase activity | GO:0016779 | 135 |
| TBP-class protein binding | GO:0017025 | 67 |
| **Aquatic Marine** | | |
| deoxyribonuclease IV (phage-T4-induced) activity | GO:0008833 | 1353 |
| sulfotransferase activity | GO:0008146 | 1232 |
| **Aquatic Freshwater** | | |
| chitinase activity | GO:0004568 | 1957 |
| glycosyltransferase activity | GO:0016757 | 1677 |
| 3'-5' exonuclease activity | GO:0008408 | 1657 |
| **Mammals** | | |
| methylenetetrahydrofolate dehydrogenase (NADP+) activity | GO:0004488 | 37 |
| dihydrofolate reductase activity | GO:0004146 | 20 |
| DNA topoisomerase activity | GO:0003916 | 20 |

| | | |
|---|---|---|
| **Engineered** | | |
| uroporphyrinogen decarboxylase activity | GO:0004853 | 33 |
| proton-transporting ATPase activity,rotational mechanism | GO:0046961 | 22 |
| **Plants** | | |
| protein dimerization activity | GO:0046983 | 352 |
| ADP binding | GO:0043531 | 230 |
| microtubule motor activity | GO:0008017 | 157 |
| copper ion binding | GO:0005507 | 132 |
| cysteine-type deubiquitinase activity | GO:0004843 | 72 |
| quinone binding | GO:0048038 | 75 |
| oxidoreductase activity,acting on NAD(P)H | GO:0016651 | 70 |
| cellulose synthase (UDP-forming) activity | GO:0016760 | 70 |
| ATP-dependent chromatin remodeler activity | GO:0140658 | 68 |
