## Supplementary material for "Quadrupling the protein family space with global metagenomics": Supplementary Table 5.pdf

**Supplementary Table 5:** Biome-specific structural domains among the most frequent, including their corresponding CATH superfamily IDs and the number of family model hits within each domain

| DOMAIN NAME | CATH SUPERFAMILY | NUMBER OF FAMILIES |
|---|---|---|
| **Aquatic Freshwater** | | |
| Metalloproteases (zincins), catalytic domain | 3.30.2010.10 | 45 |
| SGNH hydrolase | 3.40.50.1110 | 40 |
| **Aquatic Marine** | | |
| Rabenosyn, Rab binding domain | 4.10.860.20 | 33 |
| B family DNA polymerase finger domain | 1.10.287.690 | 25 |
| Conserved hypothetical protein from pyrococcus furiosus, ,ParB domain | 3.90.1530.10 | 12 |
| **Aquatic others** | | |
| Gamma-adaptin ear (GAE) domain | 2.60.40.1230 | 5 |
| Calcium-transporting_ATPase, cytoplasmic domain N | 3.40.1110.10 | 4 |
| Adaptor protein Cbl, N-terminal domain | 1.20.930.20 | 3 |
| Porphobilinogen_deaminase,_C-terminal_domain | 3.30.160.40 | 3 |
| HPT domain | 1.20.120.160 | 2 |
| Golgi alpha-mannosidase II; domain 4 | 2.70.98.30 | 2 |
| Fe,Mn superoxide dismutase (SOD) domain | 1.10.287.990 | 2 |
| Nucleotidyltransferases domain 2 | 1.20.120.330 | 2 |
| **Terrestrial** | | |
| TATA-Binding Protein | 3.30.310.160 | 55 |
| Mog1/PsbP, alpha/beta/alpha sandwich | 3.40.1000.10 | 42 |
| Alpha-catenin/vinculin-like | 1.20.120.230 | 31 |
| Dynein light chain 2a, cytoplasmic | 3.30.450.30 | 22 |
| BAG domain | 1.20.58.120 | 21 |
| STAS domain | 3.30.750.24 | 18 |
| Kinase_associated domain 1, KA1 | 3.30.310.80 | 18 |
| Membrane associated eicosanoid/glutathione metabolism-like domain | 1.20.120.550 | 18 |
| Carboxypeptidase-like, regulatory domain | 2.60.40.1120 | 16 |
| HR1 repeat | 1.10.287.160 | 15 |
| **Mammals** | | |
| EF-hand | 1.10.238.10 | 6 |
| Glycosyl_hydrolase_domain;_family_43 | 2.115.10.20 | 5 |
| Death_Domain Fas | 1.10.533.10 | 4 |
| Endonuclease/exonuclease/phosphatase | 3.60.10.10 | 4 |
| RNA polymerase II/Efflux pump adaptor protein barrel-sandwich hybrid domain | 2.40.50.100 | 4 |
| Protein_Inhibitor_Of_Neuronal_Nitric_Oxide_Synthase | 3.30.740.10 | 3 |

| | | |
|---|---|---|
| Elongation Factor G (Translational Gtpase), domain 3 | 3.30.70.870 | 3 |
| Phox-like_domain | 3.30.1520.10 | 3 |
| t-snare_proteins | 1.20.58.400 | 3 |
| Translation factors | 2.40.30.10 | 3 |
| Hypothetical_protein_af1432 | 1.10.3210.10 | 2 |
| HscB, C-terminal domain | 1.20.1280.20 | 2 |
| Metallocarboxypeptidase-like | 3.30.70.340 | 2 |
| Resolvase N-terminal catalytic domain | 3.40.50.1390 | 1 |
| **Engineered** | | |
| Release factor | 1.20.58.410 | 2 |
| Phage minor tail protein U | 3.30.70.1700 | 2 |
| **Host Others** | | |
| Lrp/AsnC effector binding domain/regulation of amino acid metabolism (RAM) domain | 3.30.70.920 | 2 |
| **Humans** | | |
| Glucose Permease (Domain_IIA) | 2.70.70.10 | 2 |
