## Supplementary figures and images for "Quadrupling the protein family space with global metagenomics"

### Supplementary_Figure_1.png

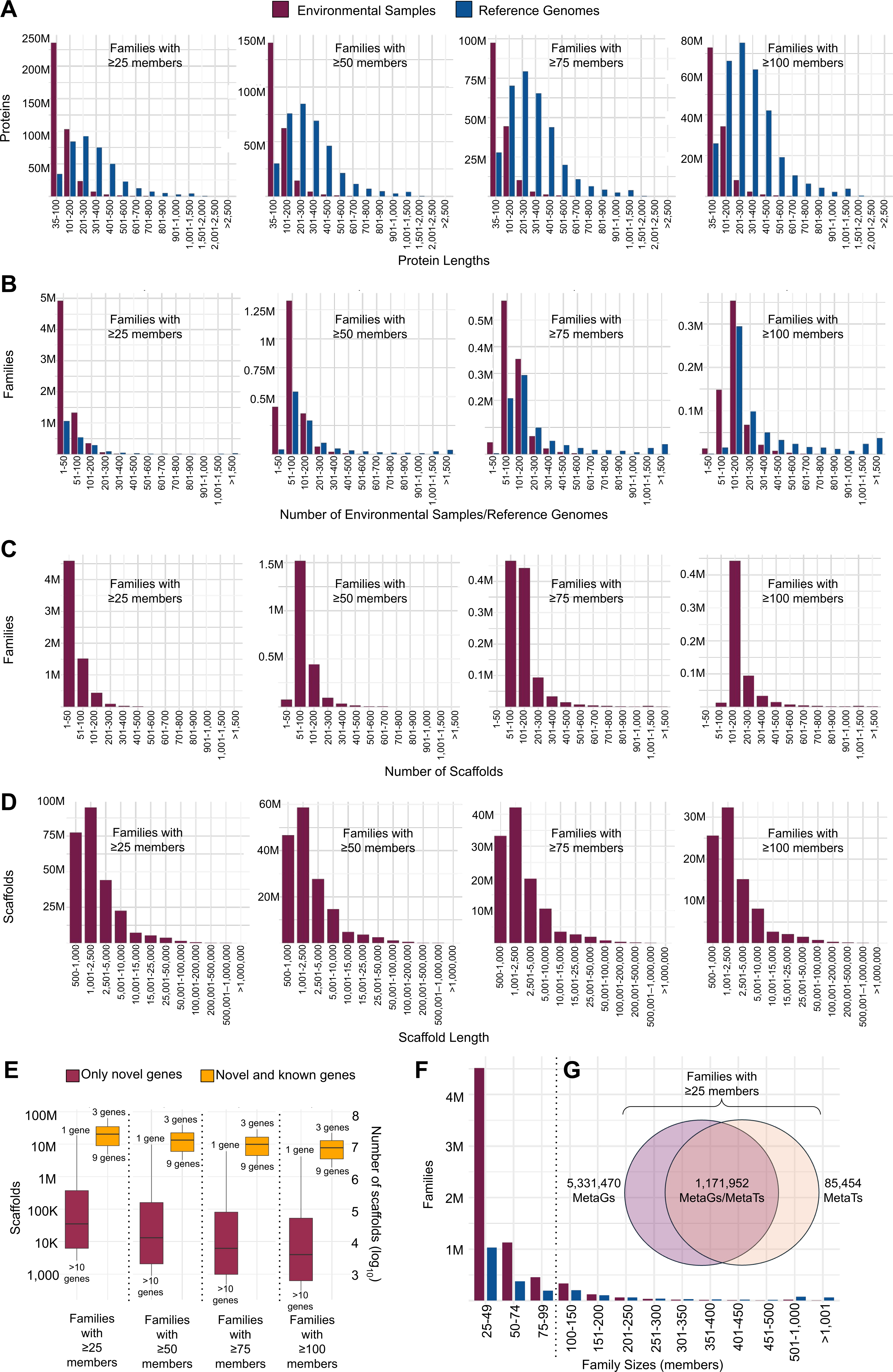

### Supplementary_Figure_2.png

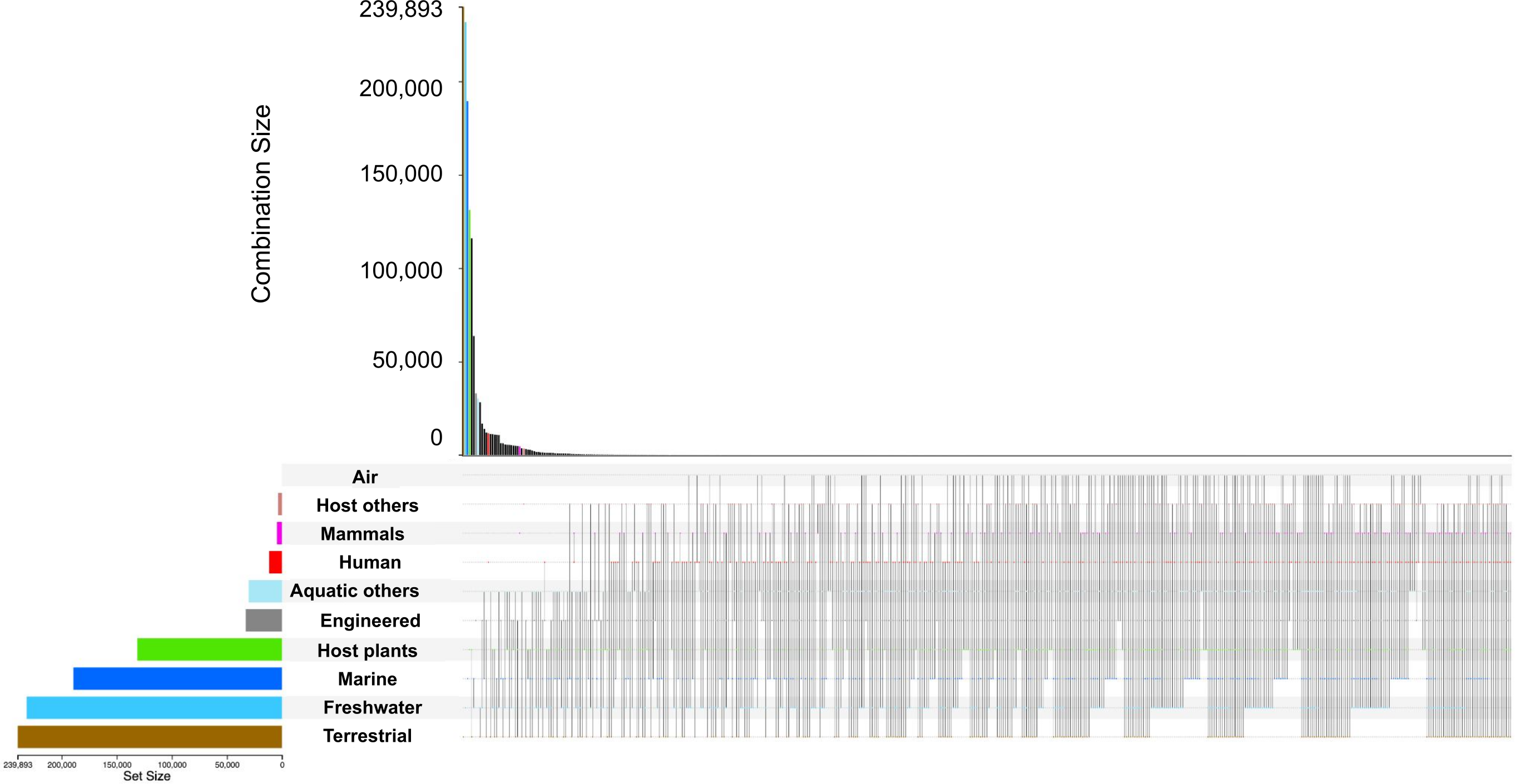

### Supplementary_Figure_3.png

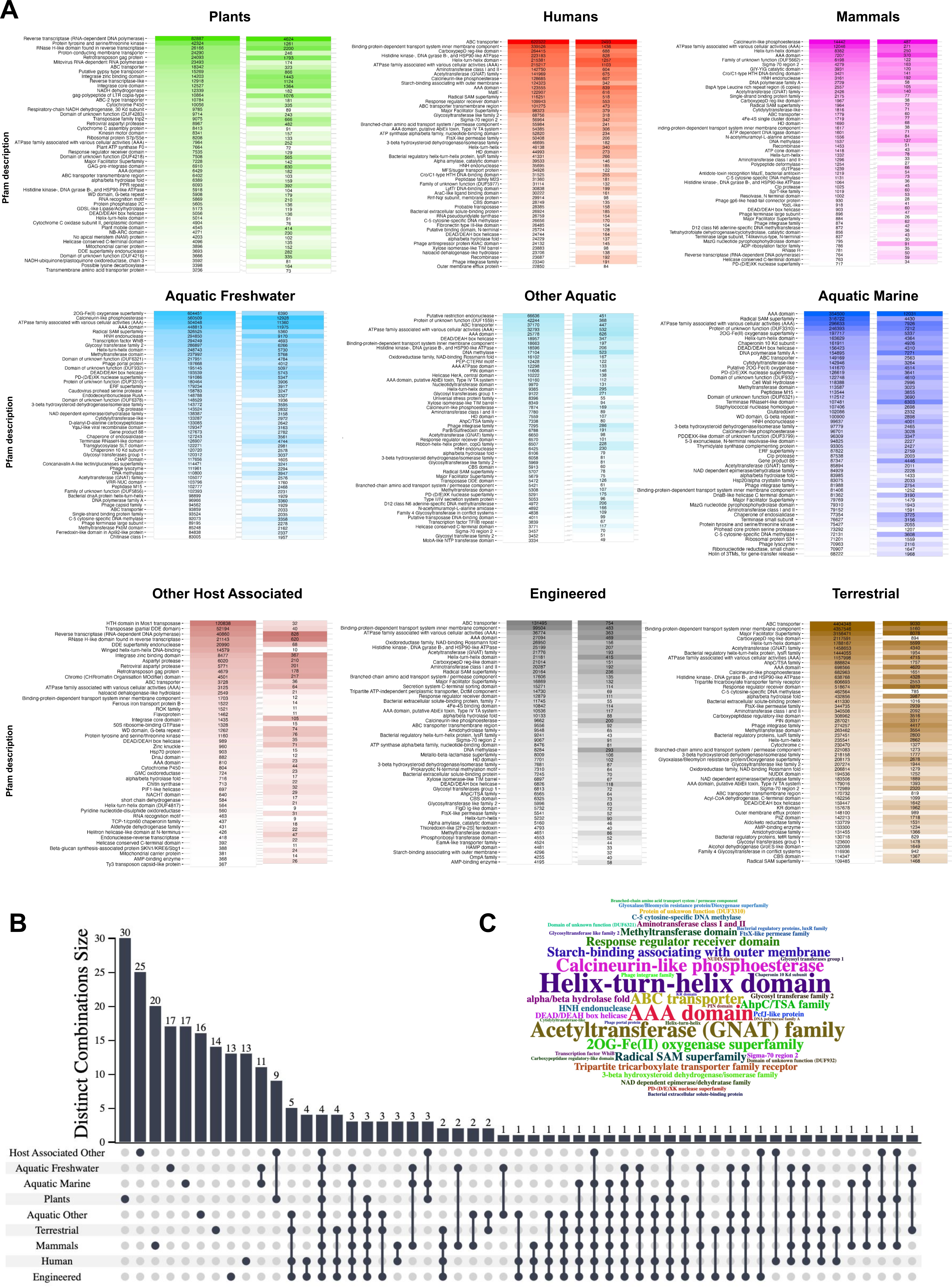

### Supplementary_Figure_4.png

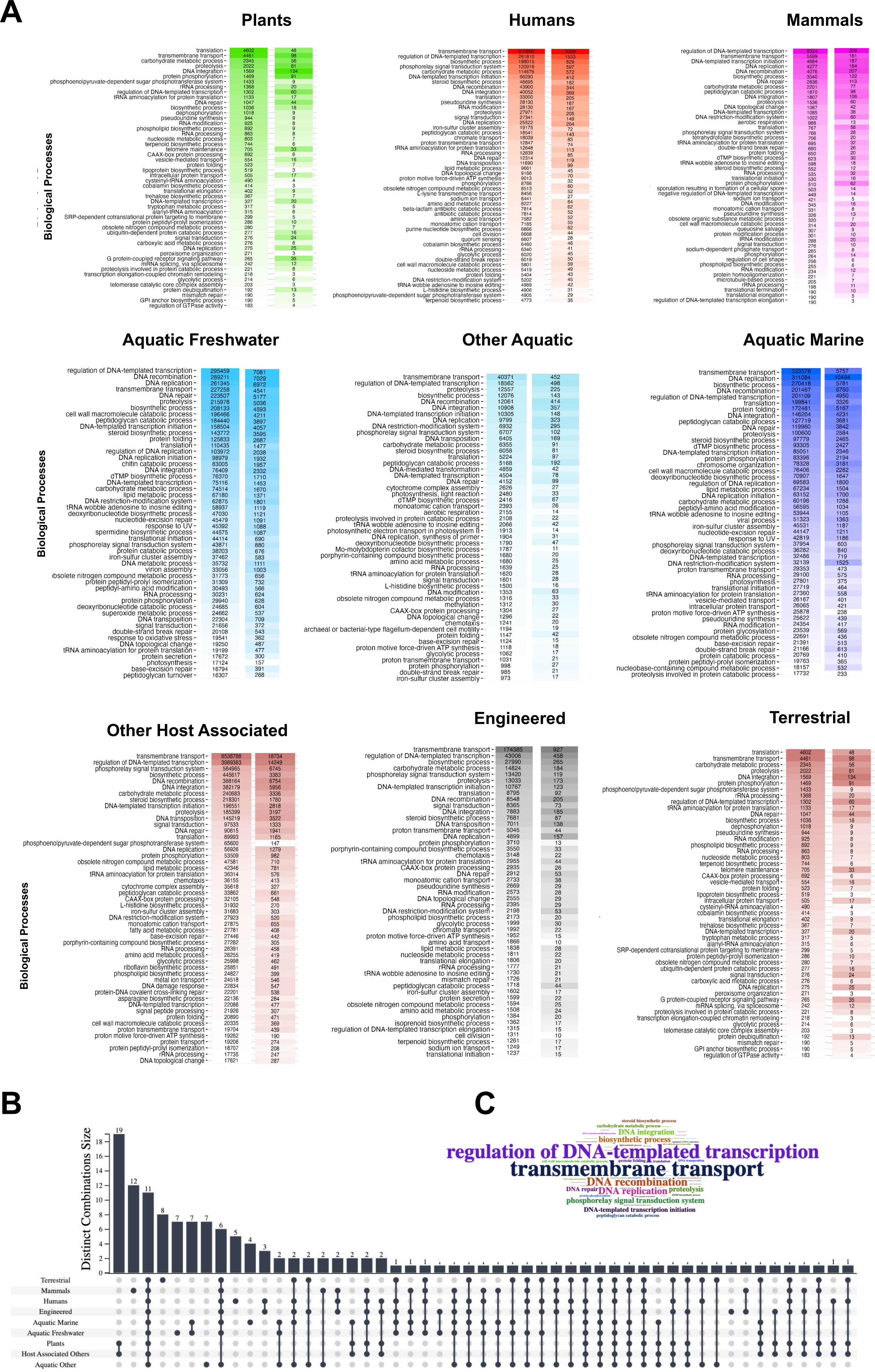

### Supplementary_Figure_5.png

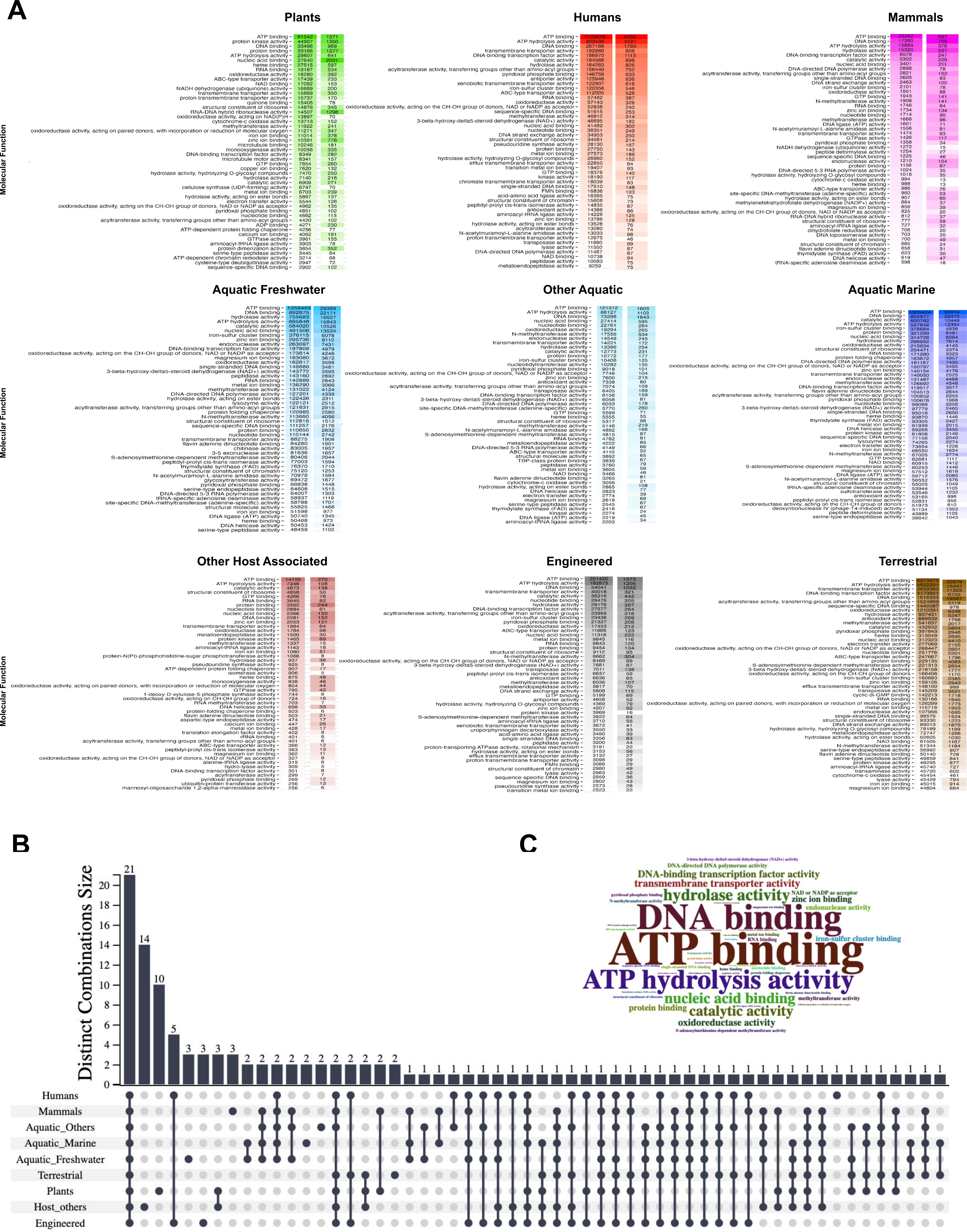

### Supplementary_Figure_6.png

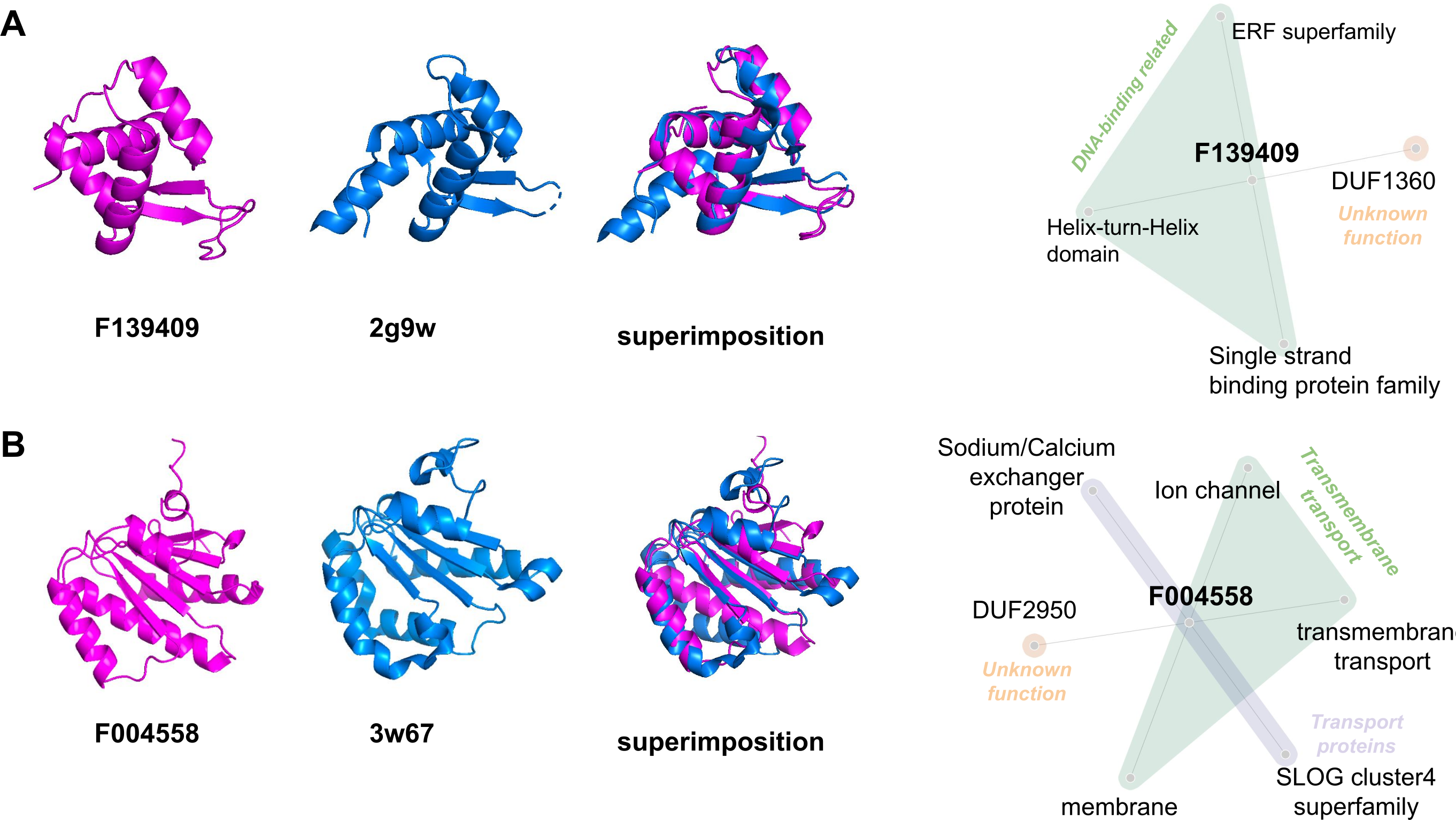
